# Subcellular multiomic profiling with SPACE-immune reveals divergent immune activation states and intracellular signaling of PBMCs

**DOI:** 10.64898/2026.11.25.769295

**Authors:** Yi Cui, Mengwei Hu, Fangfang Bai, Su Ma, Dan McGuire, Jessica McGory, Shamsudheen Karuthedath Vellarikkal, Alex Tamburino, Rebecca Mathew, Isabel Lee, Ankush Tyagi, Eric Hobbs, Michael Patrick, Shanshan He, Vanessa Peterson

**Affiliations:** Bruker Spatial Biology, Seattle, WA, USA; Merck & Co., Inc., Rahway, NJ, USA

## Abstract

Peripheral blood mononuclear cells (PBMCs) represent the most clinically accessible window into systemic immune status, yet many current single-cell technologies, such as flow cytometry and droplet-based sequencing, lack subcellular spatial resolution. Here we present SPACE-immune (SPAtial Cell Exploratory for Immune characterization), which adapts imaging-based spatial molecular profiling to measure the whole transcriptome (18,935 genes) and 64 proteins in the same PBMC at subcellular resolution. In a paired 12-donor cytokine-stimulation cohort, hierarchical multimodal cell typing resolved 17 immune subtypes. Protein-guided gating of phenotypically uniform naïve CD4+ T cells, coupled with single-cell whole-transcriptome measurements revealed two distinct transcriptional activation pathways of naïve CD4+ T cells – direct cytokine stimulation and bystander IFN-associated reprogramming, and the stimulation shifted the Treg:Th17 balance with divergent lineage-specific transcriptional responses. Uniquely, subcellular transcript localization enabled per-cell, per-gene measurement of nuclear-to-cytoplasmic RNA ratios (NCR). Using this platform, we uncover previously inaccessible biology: a novel NF-κB p65 nuclear translocation framework that links subcellular protein localization to transcriptome-wide NCR shifts, revealing cell-type-specific responses of signal-to-transcription coupling. Hence, SPACE-immune enables subcellular spatial multiomics of circulating immune cells as a blood-based platform for mechanistic immunoprofiling and translational immune monitoring.

## Introduction

Peripheral blood represents the most accessible biological material in clinical medicine, and PBMC immunoprofiling has become a cornerstone of translational research, spanning autoimmune disease monitoring, oncology biomarker discovery, and vaccine response characterization [1–5]. The dominant technologies for single-cell PBMC analysis, including droplet-based scRNA-seq [6], CITE-seq [7], CyTOF [8], and related multimodal sequencing platforms, have delivered transformative insights into immune cell diversity and function. However, these approaches share a fundamental architectural constraint: molecular information is extracted from cells in solution, inherently discarding the spatial organization of molecules within the cell. For immune cells, whose functional states are tightly regulated by the subcellular partitioning of key effectors including transcription factors, mRNAs and signaling intermediates, this is not merely a technical limitation but a biological blind spot.

Here we describe SPACE-immune, which differs from established on-slide cell-based assays (*e.g.,* image cytometry [9]) in three complementary ways. First, SPACE-immune enables significantly deeper multimodal measurement than image cytometry for immunoprofiling from real-world PBMC donors using a whole-transcriptome RNA panel (18,935 genes), paired with a high-plex immuno-oncology protein antibody panel (64 proteins) in addition to 5 morphological markers. Second, this deeper multimodal measurement enables HieraType [10], a hierarchical multimodal cell-typing algorithm that jointly integrates RNA and protein information via covariance PCA into a graph-smoothed Gaussian mixture model classification framework, resolving 17 granular PBMC subtypes with orthogonal transcriptomic and proteomic evidence. Third and most distinctively, SPACE-immune treats subcellular spatial information as a primary analytical dimension rather than metadata, such as the nuclear versus cytoplasmic compartmentalization of individual transcripts. This enables the identification of a NF-κB p65 mediated nuclear translocation framework that physically couples subcellular protein localization to transcriptome-wide NCR shifts within the same cell, a capability not available with conventional single-cell sequencing-based methods.

We applied SPACE-immune to 12 paired donors in a controlled cytokine stimulation experiment and found multiple, complementary layers of immune response that are not accessible with single-modality assays. First, naïve CD4+ T cells, as defined by protein profiling, displayed distinct transcriptional activation pathways while retaining their canonical surface protein identity, indicating a protein-RNA temporal dissociation detectable only when protein and RNA are measured in the same cell. Second, PBMC compositional analysis identified a treatment-induced shift toward Th17 dominance, including a 34% reduction in the mean Treg:Th17 ratio, alongside divergent transcriptional programs in Treg and Th17 populations. Third, the NF-κB p65 NCR framework revealed a structured hierarchy of signal-to-transcription coupling across PBMC lineages, demonstrating that protein-level pathway activation and transcriptome-wide response are dissociable properties that vary by cell type. Together, these results highlight SPACE-immune as a subcellular spatial multiomics platform for blood-based immune profiling with mechanistic depth inaccessible to prior approaches.

## Results

### A whole-transcriptome, high-plex protein spatial profiling workflow for patient-derived PBMC

We developed SPACE-immune to enable simultaneous whole-transcriptome RNA and high-plex protein profiling of suspension PBMC at subcellular resolution using CosMx^®^ SMI (**Fig. 1A**) [11]. PBMCs isolated from donor whole blood are deposited as arrayed spots onto poly-L-lysine (PLL)-coated slides, exploiting the electrostatic affinity between PLL’s multivalent polycation chains and the glycocalyx on the surface of circulating immune cells (**Fig. S1**). This surface chemistry is preferred over aminoalkylsilane-based alternatives for suspension cells due to PLL’s higher charge density and conformational flexibility and is stable under implemented mild proteinase conditions. Sequential imaging cycles first acquire a 64-plex oligo-conjugated antibody panel of immuno-oncology targets via fluorescent readout reporter probes, together with 5 morphological markers, followed by whole-transcriptome RNA detection using CosMx WTX *in situ* hybridization (ISH) probe set targeting 18,935 genes (**Fig. 1B**) [12].

**Figure 1.**
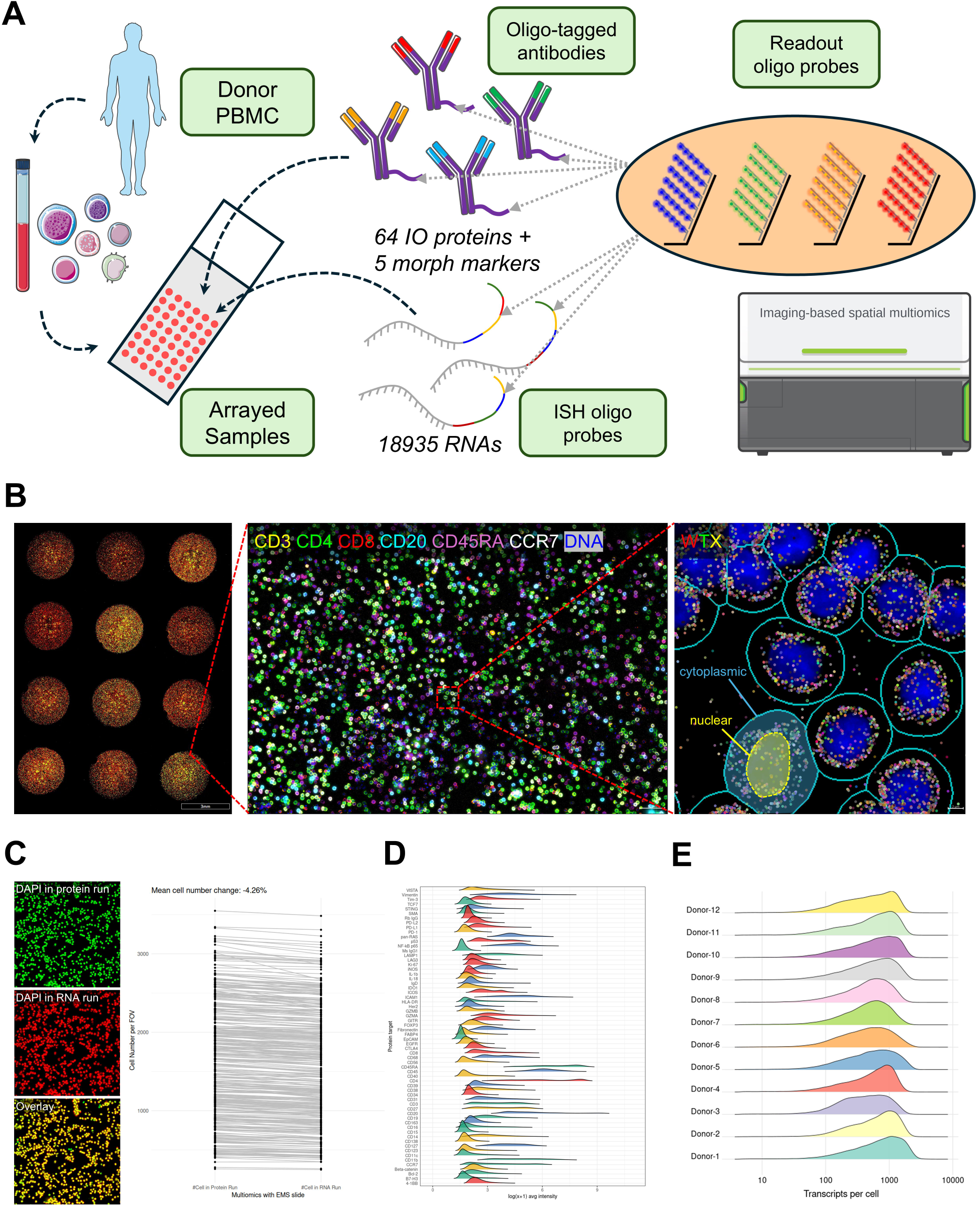
A multimodal spatial profiling workflow for simultaneous single-cell proteogenomic characterization of PBMCs. **(A)** Schematic of the SPACE-immune workflow. Peripheral blood mononuclear cells (PBMCs) isolated from donor whole blood are arrayed onto electrostatically functionalized slides in spatially indexed fields of view. Sequential CosMx SMI imaging cycles acquire protein expression using a 64-plex immunooncology (IO) antibody panel conjugated to oligonucleotide tags (plus 5 morphological markers), followed by whole-transcriptome (WTX) RNA profiling using 18,935-plex *in situ* hybridization (ISH) probe sets targeting the full protein-encoding panel. Fluorescent readout oligo probes (reporters) enable single-molecule detection with subcellular spatial resolution. **(B)** Representative composite fluorescence images illustrating multi-sample arrayed acquisition. Left: overview of a 4×3 PBMC spot array acquired in a single imaging run, demonstrating high-density spatial coverage. Center: single-cell resolved protein co-expression across the array, with simultaneous marker detection, including CD3 (yellow), CD4 (green), CD8 (red), CD20 (cyan), CD45RA (magenta), CCR7 (white), and nuclear DAPI (blue). Right: representative field-of-view showing subcellular compartment-resolved RNA transcript detection in the WTX RNA imaging run. Cell segmentation boundaries (cyan) delineate cytoplasmic (outer annular) and nuclear (inner, dashed) compartments at single-cell resolution, enabling nuclear-to-cytoplasmic transcript ratio quantification for each cell. Individual RNA molecules are visible as discrete fluorescent puncta. **(C)** Cross-run cell registration and retention quantification. Left: representative DAPI channel images from the protein imaging run (green), the RNA imaging run (red), and their overlay (yellow), demonstrating high-fidelity co-registration of sequential acquisition cycles. Right: cell count concordance across all sequentially acquired paired samples, showing a mean cell number change of −4.26% between protein and RNA runs, confirming minimal cell loss across the dual-modality workflow. **(D)** Ridgeline density plots showing the dynamic range and distribution of log-transformed mean protein expression intensities for all 64 IO panel targets (y-axis) across the PBMC cohort, stratified by condition (color). Broad dynamic range and unimodal-to-bimodal protein distributions confirm robust single-cell resolved detection across diverse surface and intracellular markers. **(E)** Distribution of per-cell total RNA transcript counts (log10-scaled) across the full PBMC array, demonstrating uniform transcriptome capture across the arrayed spot layout and confirming consistent RNA detection quality across the slide surface in the multiomic workflow.

Cell segmentation is performed independently in each imaging run using a pre-trained model, with co-registration of protein and RNA runs verified by channel-wise cross-correlation across paired FOVs. Across all samples and FOVs, the mean cell number change between protein and RNA runs was −4.26% (**Fig. 1C**), confirming minimum cell loss across the dual-modality workflow. RNA transcript detection exhibited a broad dynamic range across all 18,935 panel genes, with protein expression distributions showing unimodal-to-bimodal intensity profiles consistent with single-cell resolved detection (**Fig. 1D**). Detected RNA counts distribution confirmed uniform transcriptomic capture across the arrayed spot layout, reaching 800-1000 transcripts per cell (**Fig. 1E**), supporting unbiased cell lineage annotation (**Fig. S2**). Each detected RNA molecule is assigned to either the nuclear or cytoplasmic compartment of its parent cell (**Fig. 1B**, **right panel**), providing a per-cell, per-gene subcellular localization readout – the foundational measurement enabling the nuclear-to-cytoplasmic ratio (NCR) analyses described below.

In contrast to prior imaging-based single-cell analysis, which demonstrated PBMC profiling with limited protein markers or targeted gene panels [13, 14], SPACE-immune applies whole-transcriptome coverage as the basis for unbiased discovery, complemented by 64+ relevant protein markers. This profiling depth enables exploration of transcriptome-wide regulatory network, transcription-translation cooperation, and hypothesis-free differential expression analysis at scale – capabilities that targeted panels cannot provide. The multi-sample spot array format is flexible in the number of donors and the number of cells profiled per slide within the available imageable area, allowing for either more donors with fewer cells or fewer donors with more cells (**Fig. S7**). We used this array format to profile tens of paired donor samples – 12 control and treated pairs – within a single experiment, with spatial co-registration of protein and RNA data eliminating the need for computational deconvolution or cross-modality imputation.

### Protein-based spatial gating identifies naïve CD4+ T cell activation substates with divergent transcriptional trajectories

A key SPACE-immune capability is the use of protein expression – measured at single-cell resolution in the same physical assay as RNA – for *in silico* cell isolation, recapitulating flow cytometric gating logic within the spatial data directly (**Fig. S3**). To demonstrate this, we isolated naïve CD4+ T cells using a five-marker hierarchical protein gating strategy: CD45+ (leukocytes) → CD20−, CD3+ (T cells) → CD4+, CD8− (CD4+ T cells) → CCR7^hi,^ CD45RA+ (naïve) (**Fig. 2A-B**). This yielded a high-purity naïve CD4+ T cell population that was then subjected to whole-transcriptome sub-clustering independently of protein identity.

**Figure 2.**
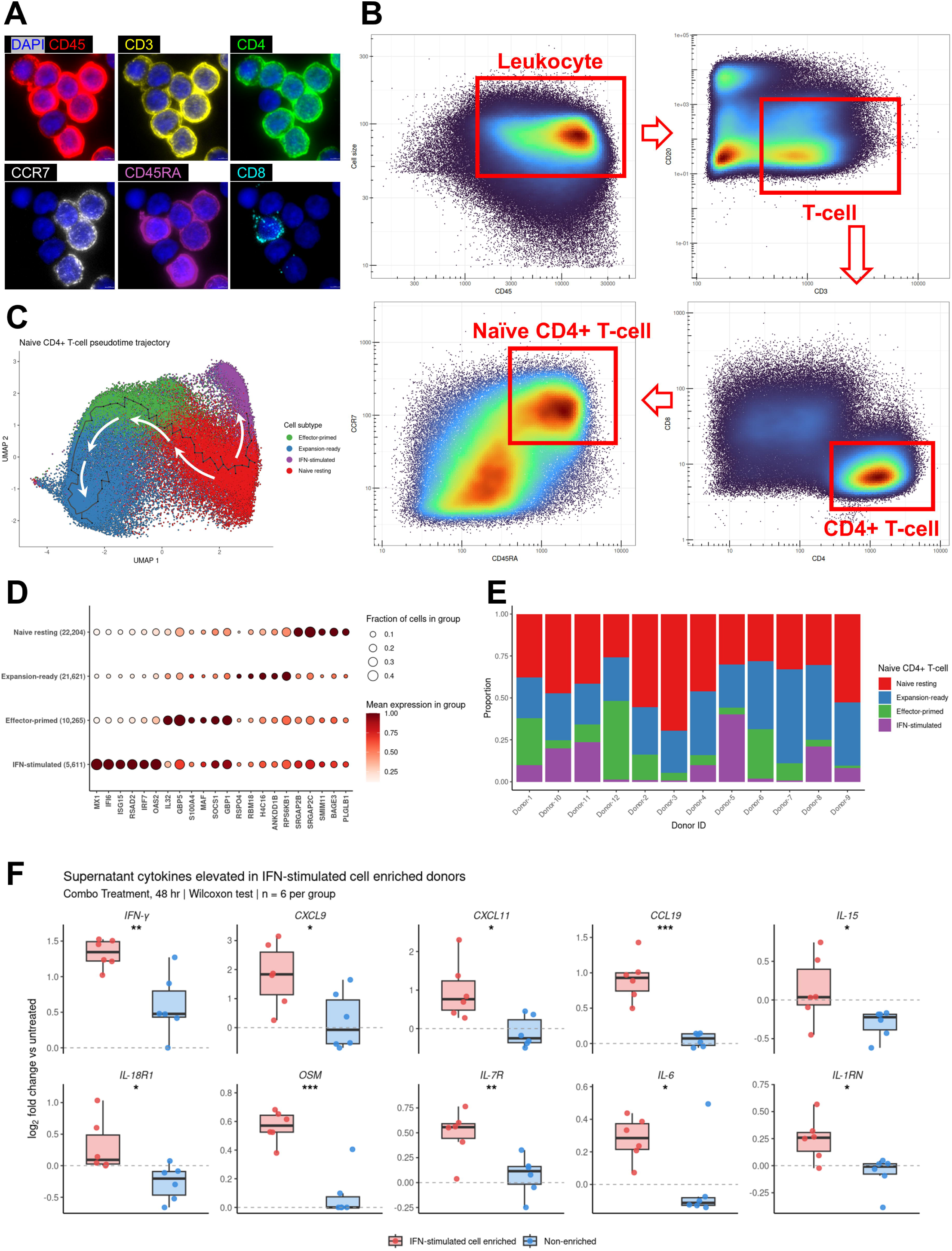
Spatial protein gating identifies naive CD4+ T cell activation substates with divergent transcriptional reprogramming. **(A)** Subcellular single-cell immunofluorescence images illustrating CosMx SMI protein channel quality for canonical immune lineage markers. Individual panels show DAPI (nuclear, blue) and CD45 (red), CD3 (yellow), CD4 (green), CCR7 (gray), CD45RA (purple), and CD8 (cyan) detected at single-cell resolution, confirming clear membrane and cytoplasmic protein localization suitable for computational gating. **(B)** Hierarchical sequential protein-based gating strategy recapitulating conventional flow cytometric immunophenotyping within CosMx spatial data. Scatter plots (analogous to flow cytometry dot plots) show sequential cell-level protein intensity gating: CD45+ leukocytes (cell size vs. CD45), CD3+ T cells (CD3 vs. CD20), CD4+ T cells (CD8 vs. CD4), and CCR7hiCD45RA+ naive CD4+ T cells, isolating a high-purity naive CD4+ T cell population for downstream transcriptomic subclustering. **(C)** UMAP embedding of naive CD4+ T cells, colored by transcriptionally defined substate. Monocle3 pseudotime trajectory (black arcs) overlaid on the UMAP reveals a continuous differentiation axis spanning four biologically distinct substates: Naive resting, Expansion-ready, Effector-primed, and IFN-stimulated, with trajectory root anchored in the Naive resting compartment. **(D)** Dot plot of discriminating transcriptional markers across the four naive CD4+ T cell substates. Dot size encodes the fraction of cells expressing each gene; color intensity encodes mean scaled expression. Cell counts per substate are indicated in parentheses. Marker genes distinguish Naive resting identity, proliferative priming (Expansion-ready: RPS6KB1, H4C16), effector reprogramming (Effector-primed: IL32, MAF), and interferon-driven stimulation (IFN-stimulated: MX1, ISG15, OAS2, IFI6). **(E)** Stacked bar charts showing per-donor proportional composition of naive CD4+ T cell substates across all 12 donors (x-axis). The IFN-stimulated substate (purple) is enriched in a subset of donors, while Naive resting (red) and Expansion-ready (blue) substates predominate across the cohort, revealing inter-donor heterogeneity in the transcriptional response trajectory. **(F)** Supernatant cytokine profiling stratified by IFN-stimulated substate enrichment. Box plots display log2 fold-change in secreted cytokine/chemokine levels (treated vs. untreated) for donors classified as IFN-stimulated cell-enriched (red, n = 6) versus non-enriched (blue, n = 6). IFN-γ, CXCL9, CXCL11, CCL19, IL-15, IL-18R1, OSM, IL-7R, IL-6, and IL-1RN are all significantly elevated in IFN-stimulated cell-enriched donors (Wilcoxon rank-sum test; *p < 0.05, **p < 0.01, ***p < 0.001), nominating IFN-γ as the primary upstream driver and CXCL9/CXCL11 as functional effector readouts of the bystander IFN response axis.

A central finding of this analysis is protein-RNA temporal dissociation. At 48 hours post-stimulation, cells across all four substates retain CCR7^hi^, CD45RA+ surface protein profiles, the canonical naïve T cell phenotype [15]. Conventional flow cytometry or multiplexed IF at this timepoint would classify these cells as phenotypically naïve and biologically quiescent. SPACE-immune reveals that the same cells have already undergone substantial transcriptional reprogramming, with four distinct transcriptional states diverging within a phenotypically homogeneous protein compartment. The four biologically coherent substates – Naïve resting, Effector-primed, Expansion-ready, and IFN-stimulated – are visualized by UMAP embedding with pseudotime trajectory overlay (**Fig. 2C**) and characterized by distinct transcriptional signatures (**Fig. 2D**). The Naïve resting substate is defined by absence of activation and interferon programs. Effector-primed cells upregulate canonical NF-κB and cytokine-response targets – notably IL32, GBP5, MAF, and SOCS1 – consistent with direct cytokine receptor engagement and early effector transcription factor induction. Expansion-ready cells are distinguished by elevated RPS6KB1 and H4 histone transcripts, and carry the highest mean pseudotime value, marking them as the most transcriptionally distant from Naïve resting and representing deep metabolic reprogramming rather than early transitional identity. The IFN-stimulated substate is defined by high-magnitude interferon-stimulated gene (ISG) expression: MX1, ISG15, OAS2, IRF7, RSAD2, and IFI6 – a transcriptional signature mechanistically distinct from the NF-κB-driven programs of the other three substates.

These substates are detectable only because protein and RNA are measured in the same cell simultaneously, without computational imputation or cross-modality alignment – a capability specifically enabled by the multimodal architecture of SPACE-immune.

Pseudotime topologies, together with substate marker identities and inter-donor prevalence data, define a two-pathway activation model (**Fig. S4**). Pathway-1 (Direct stimulation): All 12 donors show enrichment of both Effector-primed and Expansion-ready substates upon treatment, consistent with universal direct cytokine receptor engagement. The sequential pseudotime ordering Naïve resting to Effector-primed to Expansion-ready reflects progressive transcriptional commitment from early NF-κB target gene induction toward deep metabolic and translational reprogramming. Pathway-2 (Bystander IFN stimulation): The IFN-stimulated substate is enriched in only a subset of donors (**Fig. 2E**), and its early pseudotime branching from the naïve root indicates it represents an independent, parallel fate – not a downstream product of Pathway-1. Inter-donor heterogeneity in Pathway-2 enrichment reflects individual variation in innate immune responsiveness that determines whether a secondary paracrine IFN signal is generated within the PBMC culture.

To validate the biological basis of the bystander IFN pathway, we profiled cell culture supernatants from all 12 donors at 48 hours using a colocalization-by-linkage-based multiplex proteomics platform and compared cytokine levels between IFN-stimulated cell-enriched donors (n = 6) and non-enriched donors (n = 6) (**Fig. 2F**). Ten analytes were significantly elevated in IFN-stimulated cell-enriched donors: IFN-γ (p < 0.01), CXCL9 (p < 0.05), CXCL11 (p < 0.05), CCL19 (p < 0.001), IL-15 (p < 0.05), IL-18R1 (p < 0.05), OSM (p < 0.001), IL-7R (p < 0.01), IL-6 (p < 0.05), and IL-1RN (p < 0.05). IFN-γ is the dominant upstream signal – its significant and selective elevation in IFN-enriched donors directly implicates NK cell-mediated IFN-γ secretion as the paracrine driver of JAK-STAT1 activation in bystander naïve CD4+ T cells. CXCL9 and CXCL11, whose transcription is strictly dependent on active IFN-γ receptor signaling, serve as functional readouts confirming that secreted IFN-γ is biologically active in the cell culture microenvironment. IL-15 and OSM, both capable of engaging JAK-STAT pathways in T cells, may amplify ISG expression as co-stimulatory inputs.

### Multimodal HieraType cell typing resolves 17 PBMC subtypes and reveals treatment-induced Treg:Th17 imbalance

To enable unbiased, reproducible cell type annotation across the full PBMC dataset, we developed HieraType, a hierarchical multimodal cell-typing algorithm purpose-built for CosMx SMI data. HieraType begins by independently normalizing the RNA count matrix (quasi-Poisson Pearson residual normalization) and the protein expression matrix (γ-regression model Pearson residual normalization). These normalized matrices are then integrated via joint covariance PCA, projecting RNA highly variable genes, curated lineage marker genes, and protein features into a shared principal component space. A cell-cell k-nearest neighbor (kNN) similarity graph is constructed from this joint embedding, and graph-smoothed Gaussian mixture model posterior probabilities drive hierarchical classification across multiple levels (**Fig. 3A**).

**Figure 3.**
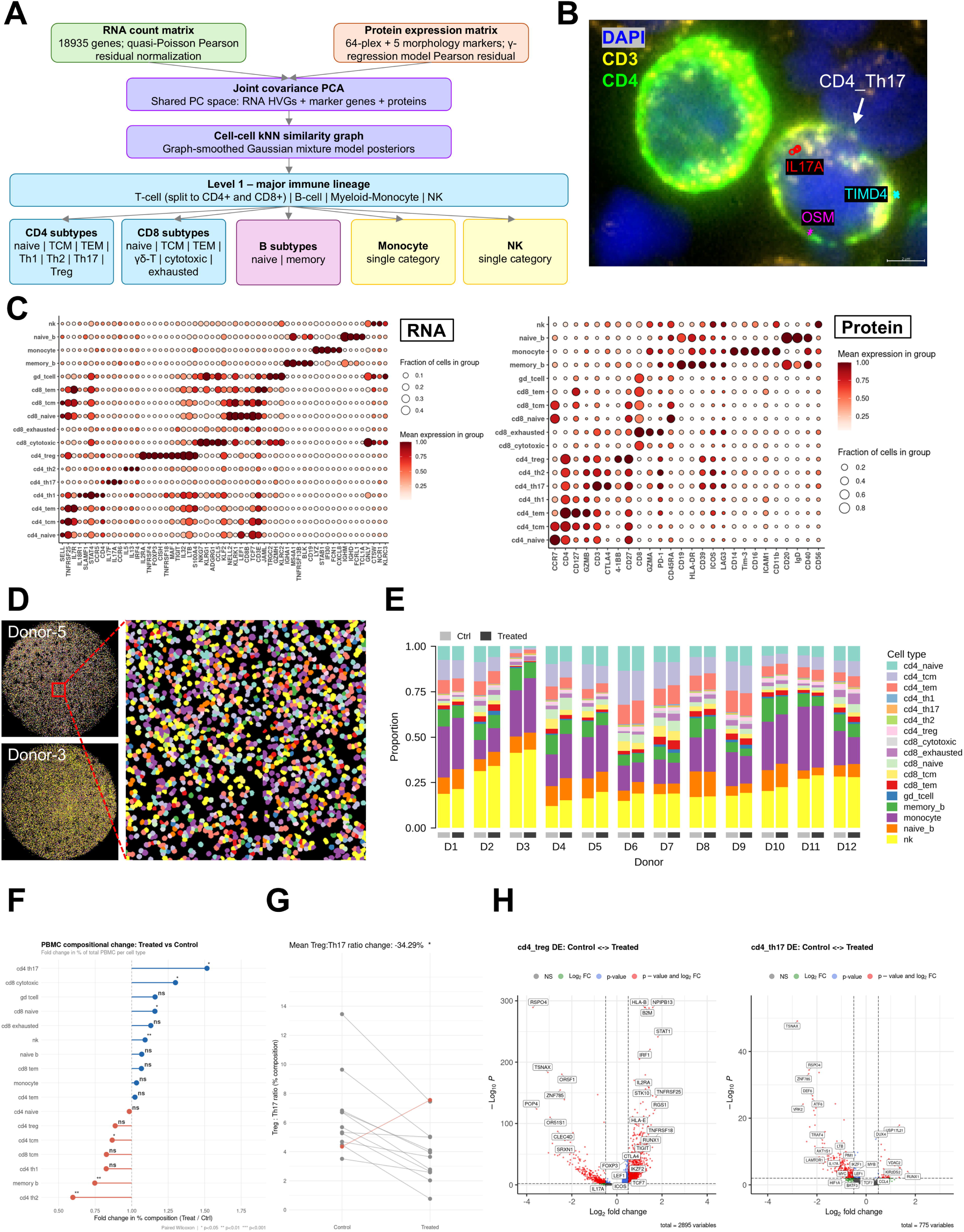
Multimodal HieraType cell taxonomy resolves 17 PBMC subtypes and uncovers a treatment-induced Treg:Th17 imbalance with lineage-specific transcriptional remodeling. **(A)** Overview of the HieraType hierarchical multimodal cell annotation pipeline. RNA count matrices (18,935 genes; quasi-Poisson Pearson residual normalized) and protein expression matrices (64-plex antibody panel plus 5 morphological markers; γ-regression model Pearson residual normalized) are jointly embedded via covariance PCA, projecting RNA highly variable genes, curated marker genes, and protein features into a shared principal component space. A cell-cell k-nearest neighbor (kNN) similarity graph is constructed and annotated by a graph-smoothed Gaussian mixture model generating posterior probability assignments. Hierarchical classification proceeds from major immune lineage (Level 1: T cell [CD4+ and CD8+], B cell, myeloid-monocyte, NK) to 17 granular subtypes across CD4, CD8, B cell, monocyte, and NK compartments. **(B)** Representative CosMx fluorescence image illustrating *in situ* validation of a CD4_Th17 cell annotation. Co-registration of DAPI (nuclear), CD3, and CD4 protein confirms T cell identity, while simultaneous single-molecule RNA detection of IL17A (Th17 effector cytokine), TIMD4, and OSM within the cell body orthogonally validates the multiomically assigned Th17 subtype identity. Scale bar, 5 μm. **(C)** Dot plots displaying canonical RNA marker gene expression (left) and protein marker expression (right) across all 17 PBMC subtypes, confirming lineage-appropriate expression patterns and discriminatory marker specificity for each annotated subtype. Dot size encodes fraction of cells expressing the marker; color intensity encodes mean scaled expression within the group. **(D)** Spatial visualization of HieraType cell type annotations *in situ*. Left panels: overview of a representative monocyte-enriched donor (top) and a NK-enriched donor (bottom) with cells pseudo-colored by assigned subtype. Right panel: corresponding zoomed fields demonstrating single-cell resolved subtype annotation across densely arrayed mixed immune populations. **(E)** Per-sample condition and per-age-group cell type compositional profiles. Stacked bar charts show the proportional abundance of all 17 subtypes across individual sample conditions, revealing sample-level variability in immune composition. **(F)** PBMC compositional change upon cytokine treatment. Lollipop plot shows the fold change in per-cell-type proportion (treated vs. control, x-axis; log-scale) across all 17 subtypes for each of 12 paired donors, with population-level significance tested by paired Wilcoxon signed-rank test. CD4_Th17, CD8 cytotoxic, and NK cells expand significantly upon treatment; CD4_Th2 and memory B cells contract. (ns: not significant; *p < 0.05, **p < 0.01, ***p < 0.001). **(G)** Paired Treg:Th17 ratio per donor (control vs. treated). Connected dot plot shows individual donor trajectories (n = 12), with a mean Treg:Th17 ratio decrease of 34.29% upon treatment (*p < 0.05, paired Wilcoxon test), indicating a shift toward Th17 immune dominance. The majority of donors (11/12) show concordant directional change. **(H)** (Left) Volcano plot of differential expression in CD4_Treg cells (treated vs. control; n = 2,895 variables). Points are colored by significance category (log2FC only, p-value only, or both). Upregulated genes include canonical Treg functional activation markers (FOXP3, IL2RA, CTLA4, TIGIT, IKZF2, TNFRSF25/DR3, TNFRSF18/GITR), while MHC class I and interferon-response genes (HLA-B, HLA-E, B2M, STAT1, IRF1) are among the most significantly upregulated transcripts, together defining a state of robust Treg functional activation. (Right) Volcano plot of differential expression in CD4_Th17 cells (treated vs. control; n = 775 variables) reflects an attenuated transcriptional response consistent with post-expansion quiescence at 48 hours. The most significantly downregulated genes include TSNAX, mTORC1 components (AKT1S1, LAMTOR1), the Th17 lineage transcription factor BATF, and IL17A itself – indicating that despite Th17 numeric expansion at the population level, per-cell effector transcription is suppressed, a dissociation only resolvable at single-cell resolution.

Primary level classification assigns four major immune lineages: T cell (further split into CD4+ and CD8+ branches), B cell, myeloid-monocyte, and NK cell. Secondary level classification resolves 17 granular subtypes: within CD4+ T cells (naïve, TCM, TEM, Th1, Th2, Th17, Treg), within CD8+ T cells (naïve, TCM, TEM, γδ-T, cytotoxic, exhausted), within B cells (naïve, memory), plus monocyte and NK as single categories. Subtype assignments were validated by orthogonal marker expression: RNA-level marker genes confirmed canonical lineage transcription factors and effector genes, while protein-level marker plots confirmed concordant surface protein expression for all subtypes (**Fig. 3C**). *In situ* spatial validation confirmed single-cell resolved subtype annotation – for example, a CD4_Th17 cell was validated by co-detection of CD3/CD4 protein with IL17A, TIMD4, and OSM RNA transcripts within the same cell boundary (**Fig. 3B**).

The cell type annotations were visualized *in situ* across the full PBMC spot array and representative zoomed fields of view, confirming single-cell resolved subtype assignment within densely packed, mixed immune populations (**Fig. 3D**). Stacked compositional profiles across all 24 sample-condition pairs revealed substantial inter-donor variability in baseline immune composition, with naïve CD4+ T cells, monocytes, NK cells, and memory_B cells showing the widest donor-to-donor range (**Fig. 3E**).

Stratification by donor age group revealed a directional shift in immune composition consistent with immunosenescence: younger donors were enriched for adaptive immune cells (*e.g.,* CD4/CD8 T-cell), while older donors showed progressive enrichment of innate lineage cells (NK, monocyte) and contraction of the naïve T cell compartment (**Fig. S5**). This age-associated compositional drift – a well-described feature of human immune aging reflecting thymic involution and homeostatic contraction of naïve T cell pools [16–18] – is directly quantifiable at 17-subtype resolution from a single SPACE-immune run, without separate immunophenotyping experiments.

Compositional analysis across 12 paired donors (control vs. treated) using paired Wilcoxon signed-rank tests identified significant population-level shifts: NK cells (+1.50×, p = 0.001), CD4_Th17 (+2.03×, p = 0.016), and CD8_cytotoxic cells (+1.76×, p = 0.034) expanded upon treatment, while CD4_Th2 (p < 0.001) and memory_B cells contracted (**Fig. 3F**). These directional changes are biologically coherent with the known biology of the cytokine stimulation protocol employed, while the specific signaling mechanisms are not elaborated here given their dependence on the proprietary stimulus composition (an IL2-involved combo).

The Treg:Th17 ratio – a key index of inflammatory balance with direct clinical relevance in autoimmunity, oncology, and transplant immunology – shifted significantly toward Th17 dominance upon treatment (mean ratio −34.29%, paired Wilcoxon p = 0.009), with 11/12 donors showing concordant directional change (**Fig. 3G**). This ratio shift is driven primarily by Th17 compositional expansion rather than Treg numeric loss. Differential expression (DE) analysis reveals that the two populations are in opposing transcriptional states: Treg cells upregulate a robust functional activation signature (FOXP3, IL2RA/CD25, CTLA4, TIGIT, IKZF2/Helios, TNFRSF25/DR3, TNFRSF18/GITR), while Th17 cells show a post-activation quiescence signature characterized by downregulation of IL17A, stemness factors (LEF1, TCF7), and mTOR pathway components (AKT1S1, LAMTOR1, PIM1) (**Fig. 3H**). This observed paradox – simultaneous Treg functional activation and Th17 numeric expansion – indicates that the treatment drives both pro-inflammatory effector amplification and an attempted counter-regulatory response, with the net immunological balance tipping toward Th17 dominance at 48 hours. Mirroring the Treg pattern, CD4_Th1 cells similarly showed transcriptional activation without numeric expansion: a strong activation-dominant DE profile (STAT4, RUNX1, BATF, TNF upregulated; 1,302 vs. 217 genes up vs. down) in the absence of significant compositional change (FC = 0.826, p = 0.077), indicating that cytokine stimulation drives effector priming across helper T cell lineages rather than selective clonal expansion (**Fig. S6**). Donor-9 was identified as a directional outlier for the Treg:Th17 ratio, retaining an elevated ratio post-treatment, warranting further investigation into inter-individual variation in regulatory versus effector T cell responsiveness.

### A nuclear-to-cytoplasmic ratio framework couples NF-κB p65 subcellular localization to transcriptome-wide compartmentalization

The defining analytical capability of imaging-based suspension cell profiling – one absent from conventional sequencing-based PBMC technologies – is the ability to measure the subcellular distribution of individual molecules within each cell. SPACE-immune assigns each detected RNA transcript to either the nuclear or cytoplasmic compartment of its parent cell via segmentation of DAPI-defined nuclear boundaries (**Fig. 4A**, **lower**). We extended this to protein by measuring NF-κB p65 nuclear-to-cytoplasmic intensity ratio (NCR) at single-cell resolution, enabling a physically grounded readout of canonical NF-κB pathway activation state.

**Figure 4.**
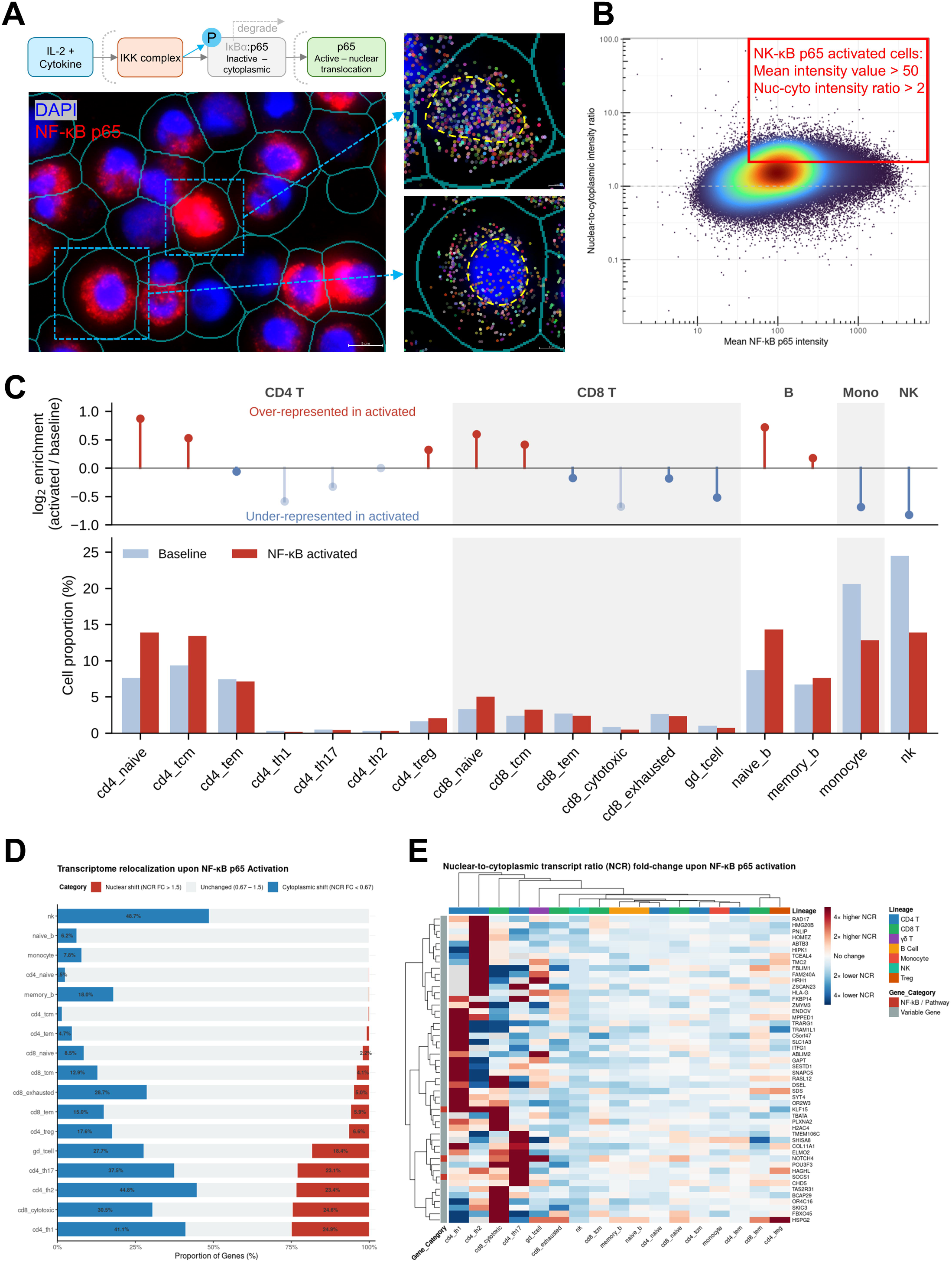
NF-κB p65 nuclear translocation as a spatially resolved single-cell activation readout links subcellular protein dynamics to cell-type-specific transcriptional reprogramming. **(A)** Schematic and imaging basis of NF-κB p65 nuclear-to-cytoplasmic ratio (NCR) quantification. Top: canonical NF-κB signaling pathway in which upstream activating signals (exemplified by IL-2 cytokine) engage the IKK complex, driving IκBα phosphorylation and degradation, releasing p65 for nuclear translocation and transcriptional activation; in the resting state, p65 is sequestered as an inactive cytoplasmic IκBα:p65 complex. Bottom: representative CosMx SMI image showing simultaneous DAPI (blue, nuclear) and NF-κB p65 (red) immunofluorescence with segmentation boundaries (cyan) overlaid, enabling single-cell nuclear and cytoplasmic compartment-resolved p65 intensity quantification. Insets illustrate a p65-activated cell (nuclear-enriched, yellow dashed boundary) and an inactivated cell (cytoplasmic, blue nucleus) at high magnification. **(B)** Two-dimensional bi-axial gating strategy for NF-κB p65 activated cell classification. Scatter plot of mean NF-κB p65 fluorescence intensity (x-axis, log10) versus nuclear-to-cytoplasmic intensity ratio (y-axis, log10) across all cells in the cohort. Activated cells (upper-right gate, red box) are defined by dual criteria: mean p65 intensity > 50 a.u. and nuclear-to-cytoplasmic ratio > 2, capturing cells with both sufficient p65 abundance and confirmed nuclear enrichment. **(C)** Cell-type composition of NF-κB p65 baseline and activated populations. Bottom: percentage of all baseline (light blue) and all NF-κB-activated (red) cells assigned to each subtype; each color sums to 100%. Top: log2 enrichment of each subtype among activated relative to baseline cells; faded points denote subtypes contributing <1% to both populations. Shaded bands indicate major lineages. **(D)** Transcriptome-wide nuclear-to-cytoplasmic transcript relocalization upon NF-κB p65 activation, stratified by cell type. Horizontal stacked bar chart displays the proportion of genes (x-axis) in each subtype classified as nuclear-shifted (NCR fold-change > 1.5; red), unchanged (0.67–1.5; gray), or cytoplasmic-shifted (NCR fold-change < 0.67; blue) in NF-κB activated versus baseline cells. CD4_Th1 and CD8 cytotoxic cells exhibit the highest proportions of cytoplasmic-shifted transcripts, consistent with negative feedback-driven IκBα resynthesis and NF-κB target gene export; NK cells exhibit predominant cytoplasmic transcript redistribution reflecting a distinct translational response architecture. **(E)** Biclustered heatmap of NCR fold-change for NF-κB pathway genes and variable genes across all 17 PBMC subtypes (columns) and selected gene features (rows), in NF-κB activated versus baseline cells. Gene categories (NF-κB pathway, color-coded; variable genes, gray) and immune lineage (column annotation bar) are indicated. Hierarchical clustering reveals cell-type-specific patterns of nuclear transcript enrichment and cytoplasmic shift.

NF-κB p65 activation was defined by a two-gate strategy: mean p65 fluorescence intensity (excluding p65-negative/low cells) AND nuclear-to-cytoplasmic intensity ratio (confirming nuclear enrichment). This biaxial gate identified activated cells across all 17 PBMC subtypes (**Fig. 4B**). Because lineage abundance differs widely across PBMC, we compared the cell-type composition of the activated population with that of the baseline population. Activated cells were enriched for naïve and central memory lymphocytes relative to baseline: CD4_naïve (13.9% vs. 7.6%, 1.8-fold), naïve_B (14.3% vs. 8.7%, 1.6-fold), CD8_naïve (5.0% vs. 3.3%, 1.5-fold), CD4_TCM (13.4% vs. 9.3%, 1.4-fold), and CD8_TCM (3.2% vs. 2.4%, 1.3-fold). In contrast, innate lineages were under-represented among activated cells: NK (13.9% vs. 24.5%, 0.57-fold) and monocytes (12.8% vs. 20.6%, 0.62-fold). CD4_TEM, CD8_TEM, CD8_exhausted, CD4_Treg, and memory_B showed comparable representation in both populations (within 0.9-1.3-fold), whereas CD4_Th1, Th2, Th17, CD8_cytotoxic, and γδ T cells each contributed <1% of either population (**Fig. 4C**).

To couple protein-level p65 translocation to transcriptional output, we computed per-gene NCR fold-change (NCR FC) in NF-κB p65-activated versus baseline cells for each cell type, classifying genes as nuclear-shifted (NCR FC > 1.5), cytoplasmic-shifted (NCR FC < 0.67), or unchanged (0.67-1.5) (**Fig. 4D**). Cytoplasmic-shifted genes outnumbered nuclear-shifted genes in every cell type, but the extent and directionality of relocalization varied markedly. NK cells showed the most extensive unidirectional redistribution (48.7% of genes cytoplasmic-shifted, with negligible nuclear shift). CD4_Th1, CD4_Th2, CD4_Th17, CD8_cytotoxic, and γδ T cells showed bidirectional redistribution, combining large cytoplasmic-shifted fractions (27.7-44.8%) with substantial nuclear-shifted fractions (18.4-24.9%). CD4_Treg, CD8_TEM, CD8_TCM, CD8_exhausted, and memory_B showed intermediate, predominantly cytoplasmic shifts (12.9-28.7% cytoplasmic; ≤6.6% nuclear), whereas naïve and central memory lymphocytes (CD4_naïve, CD4_TCM, CD8_naïve, naïve_B), CD4_TEM, and monocytes showed minimal relocalization (≤8.5% cytoplasmic; ≤2.2% nuclear). Notably, the populations most over-represented among p65-activated cells (as in **Fig. 4C**) were those with the least transcript relocalization, indicating that p65 nuclear translocation and transcriptome-wide RNA redistribution are dissociable.

Biclustered heatmap analysis of NCR FC across NF-κB pathway genes and other variable genes for all 17 cell types reveals the gene-level architecture underlying these cell-type-specific patterns (**Fig. 4E**). Column-wise hierarchical clustering separates cell types into two broad groups: a left cluster comprising CD4_Th1, CD8_cytotoxic, CD4_Th17, CD4_Th2, and γδ_T, which show strong nuclear enrichment (deep red) in an upper gene block (including RAD17, HMG20B, PNLIP, HOMEZ, ABTB3, HBP1, TCEA4, TMC2, FBLIM1, FAM244A, and HRH1); and a right cluster encompassing NK, CD8_exhausted, memory_B, naïve_B, CD8_naïve, CD8_TCM, monocyte, CD4_TEM, CD4_naïve, CD4_TCM, and CD4_Treg, which show predominantly cytoplasmic shift (blue) or near-neutral values across the same gene set. NK cells stand out within the right cluster as having near-uniform cytoplasmic redistribution across virtually all gene features, both NF-κB pathway genes (orange annotation) and variable genes (gray), consistent with the global cytoplasmic dominance observed in **Fig. 4D**. NF-κB pathway genes (orange) do not cluster separately from variable genes at the row level, indicating that NF-κB targets are subject to the same cell-type-specific subcellular redistribution logic as the broader transcriptome – rather than exhibiting a stereotyped nuclear enrichment pattern expected from canonical pathway activation. This finding suggests that post-transcriptional regulation of mRNA localization (nuclear retention vs. cytoplasmic export) is a cell-type-specific modulator of effective NF-κB output that is only measurable by subcellular spatial multiomics.

This NF-κB p65 NCR framework constitutes a novel analytical approach for imaging-based suspension cell profiling, physically coupling subcellular protein localization to transcriptome-wide RNA compartmentalization in the same cell. Conventional single-cell sequencing-based multimodal platforms cannot provide this measurement, as sequencing inherently loses subcellular spatial information during cell lysis. STAMP, while operating on the same CosMx SMI platform [14], does not report compartmental (*e.g.,* NCR-based) analyses in its demonstration experiments, treating RNA and protein signals as whole-cell aggregates. The NF-κB p65 NCR framework thus represents the clearest demonstration of the unique biological dimension that subcellular spatial multiomics contributes to immune cell profiling.

## Discussion

SPACE-immune demonstrates that subcellular spatial multiomics of circulating immune cells is not merely a technical extension of imaging-based tissue profiling – it is a qualitatively different analytical paradigm that accesses biological dimensions invisible to all prior PBMC profiling technologies. Three findings from this study illustrate why subcellular multiomics matters for immunology.

The first is the protein-RNA temporal dissociation in naïve CD4+ T cells. At 48 hours post-stimulation, phenotypically uniform CCR7^hi^ CD45RA+ cells harbor four transcriptionally distinct activation substates, two of which (IFN-stimulated and Effector-primed) represent biologically distinct responses to different upstream signals. Flow cytometry-based or affinity-based immunophenotyping at 48 hours would classify these cells as naïve and biologically quiescent, leading to systematic mischaracterization of early immune activation. However, because SPACE-immune measures both protein and RNA in the same cell, where both signals are physically co-registered rather than computationally imputed, this dissociation can be directly observed. The two-pathway model, direct cytokine stimulation driving Effector-primed and Expansion-ready states universally, versus bystander IFN imprinting driving the IFN-stimulated state in some donors, also reveals that inter-individual variation in innate immune responsiveness creates a secondary source of naïve CD4+ T cell heterogeneity invisible to conventional flow-based phenotypic profiling. The supernatant IFN-γ data, showing significant elevation specifically in donors with enriched IFN-stimulated CD4+ T cells, validates the bystander pathway at the protein level and identifies CXCL9/CXCL11 as functional readout biomarkers of active IFN-γ receptor engagement.

The second finding is that compositional and transcriptional readouts of the same CD4+ T cell lineages can diverge. Treatment shifted the Treg:Th17 balance toward Th17 dominance, driven by Th17 expansion rather than Treg loss, yet the transcriptional states of the two lineages did not mirror these numeric changes: Treg cells engaged a functional activation program without expanding, whereas expanded Th17 cells downregulated IL17A, consistent with post-activation quiescence. A cell-type composition readout, such as flow cytometric enumeration, would report a pro-inflammatory shift; a transcriptome-only readout would emphasize counter-regulatory Treg activation. Neither readout alone would capture the net state in which effector expansion and attempted regulatory compensation coexist. Resolving this required multimodal typing of rare CD4+ T cell subsets at cohort scale and measuring abundance and transcriptional state in the same assay. Because the Treg:Th17 balance is a widely used index of inflammatory status in autoimmunity, transplantation and cancer immunotherapy, such a two-axis readout may prove more informative for immune monitoring than single-modality measures. Whether Th17 expansion at this early timepoint reflects proliferation or acquisition of Th17 transcriptional identity by pre-existing CD4+ T cells remains to be determined.

The third finding – the NF-κB p65 NCR framework – is the most technically novel contribution of SPACE-immune. By measuring nuclear-to-cytoplasmic p65 protein intensity and nuclear-to-cytoplasmic transcript ratio shifts in the same cell, SPACE-immune creates a continuous, single-cell readout of signal-to-transcription coupling that is not obtained from conventional single-cell sequencing-based approaches. The cell-type-specific patterns of transcript redistribution upon p65 nuclear translocation span three empirically distinct profiles: global cytoplasmic dominance in NK cells, bimodal redistribution in CD4 Th1 and CD8 cytotoxic cells reflecting simultaneous transcriptional induction and IκBα-driven transcript export, and predominantly cytoplasmic redistribution with minimal nuclear enrichment in monocytes and CD8 memory subsets – none of which are accessible to protein-only or RNA-only measurement, and each of which carries distinct mechanistic implications for NF-κB pathway output that are only resolvable by subcellular spatial multiomics.

Several limitations of the current implementation merit consideration. The use of a single stimulation timepoint (48 hours post-stimulation) means that the observed protein-RNA temporal dissociation and transcriptional substate distributions represent a snapshot of early-to-mid immune activation; the precise kinetics of divergence between protein phenotype and transcriptional identity, and whether additional substates emerge at later timepoints, remains to be established. Similarly, the 12-donor cohort was sized for platform demonstration and paired statistical validation rather than for clinical biomarker discovery; findings related to inter-donor heterogeneity – particularly the IFN-stimulated naïve CD4+ T-cell enrichment pattern and the Treg:Th17 change trend – should be considered preliminary pending validation in larger, more diverse real-world cohorts. The NF-κB p65 NCR framework as implemented uses a binary activation gate; future iterations incorporating continuous NCR quantification along pseudotime trajectories could resolve the temporal dynamics of signal-to-transcription coupling at even higher resolution. Finally, SPACE-immune’s sequential protein-then-RNA imaging workflow requires multi-day instrument time per run, in contrast to flow cytometry’s same-day turnaround for immunophenotyping. This tradeoff is inherent to the depth of measurement: SPACE-immune is designed for mechanistic discovery and whole-transcriptome characterization rather than as a replacement for flow cytometry, and users should select the appropriate modality based on the biological question of interest.

More broadly, SPACE-immune establishes that the diagnostic and research value of spatial transcriptomics extends beyond tissue architecture to the intracellular organization of single cells. As imaging-based platforms expand toward higher multiplex protein panels, faster cycling chemistries, and improved cell segmentation algorithms, the ability to measure where molecules are within a cell – not just how many – will become an increasingly powerful dimension of single-cell immunology. SPACE-immune demonstrates that this dimension is accessible from a standard blood draw, positioning subcellular spatial multiomics as a scalable, liquid-biopsy-compatible platform for mechanistic immune monitoring in both research and clinical contexts.

## Methods and Materials

### PBMC isolation and treatment

PBMCs were isolated from the whole blood of 12 healthy donors by density gradient centrifugation. Cells were treated with a proprietary cytokine stimulation protocol or vehicle control for 48 hours. After that, cells were fixed in 4% PFA (Electron Microscopy Sciences) at 4°C overnight. The next day, cells were washed with PBS to remove PFA, and then resuspended in PBS with 10% DMSO and slowly cooled down in a Styrofoam box in -80°C. Prior to cell deposition onto glass slides, cells were quickly thawed in a 37°C water bath, spun down and resuspended in PBS to remove the DMSO.

### Cell deposition and slide preparation

Fixed PBMCs were deposited onto fresh, dust-free poly-L-lysine (PLL)-coated glass slides (Electron Microscopy Sciences, cat. #63410-02) and dried at 42°C for 1.5–3 hours, with the precise drying duration scaled to the deposited volume to ensure complete and uniform evaporation without over-drying. To enable multiplexed profiling of tens of samples on a single slide – a critical throughput advantage for cohort-scale studies – a multi-well arrayed configuration was established using on-slide silicone isolators (*e.g.,* Grace Bio-Labs reusable gaskets), which physically partition the slide surface into discrete, leak-proof wells while remaining compatible with downstream CosMx SMI imaging workflows (**Fig. 1B**, **S7**).

We specifically selected EMS PLL-coated slides rather than pre-charged commercial alternatives such as VWR SuperFrost Plus or Leica BOND slides. Although these latter formats are widely used in histological workflows, their permanently positive surface charge renders additional PLL or poly-D-lysine (PDL) coating counterproductive: electrostatic repulsion between the pre-existing polycation layer and any exogenously applied coating interferes with uniform cell adhesion and introduces surface chemistry variability that is difficult to control across experiments. PLL-coated EMS slides, by contrast, provide a defined, reproducible adhesion substrate with a well-characterized surface charge density amenable to standardization.

The drying step requires particular attention during initial protocol optimization. We recommend closely monitoring the drying process for the first several experiments to calibrate the appropriate duration for a given deposition volume and laboratory humidity conditions, as over-drying can compromise cell morphology and RNA quality, while under-drying leads to inadequate adhesion. As a general guideline, once all visible liquid has evaporated from the slide surface, slides should be allowed to continue air-drying for an additional ∼1 hour to ensure complete removal of residual moisture from the cell monolayer. Following the drying process, slides were transferred to 70% ethanol at 4°C for storage for up to one month, providing flexibility in experimental scheduling and enabling batch processing across multiple collection timepoints.

### SPACE-immune multimodal assay

To implement the SPACE-immune multimodal assay, we developed a sequential protein-first, RNA-second readout workflow on the CosMx SMI platform, incorporating targeted modifications to the manufacturer’s default single-analyte protocols to preserve cell integrity and enable high-quality joint profiling from a single slide.

For protein sample preparation, a mild antigen retrieval step was applied at 80°C for 5 minutes – in contrast to the more stringent conditions used in standard tissue-based protocols – to prevent detachment or loss of immature and fragile cell populations, including naïve lymphocytes and progenitor-like monocytes. Slides were then incubated with the 64-plex CosMx Immuno-Oncology (IO) antibody panel, comprising antibodies targeting key immunophenotyping markers across major PBMC lineages, along with fluorescent fiducial beads for image registration and spatial alignment across imaging cycles. To fix antibody-antigen complexes in place and stabilize cells on the slide surface prior to multi-cycle imaging, slides were treated with 1 mg/mL Dithio-bis(succinimidyl propionate) (DSP), a cleavable amine-reactive crosslinker chosen for its compatibility with downstream chemical reversal. The 64 protein targets, together with 5 upfront morphological markers used for cell segmentation and quality control, were imaged using a 16-cycle configuration with five laser excitation wavelengths: 385 nm, 488 nm, 530 nm, 590 nm, and 647 nm.

Following completion of the protein imaging run, flow cells were removed and slides were treated with 25 mM Tris(2-carboxyethyl)phosphine (TCEP), a reducing agent that cleaves the disulfide bond within the DSP crosslinker, reversing antibody fixation without compromising RNA integrity. A second heat-mediated retrieval step (80°C, 5 min) was then applied to fully dissociate remaining bound IO panel antibodies, ensuring that residual protein probes did not interfere with subsequent RNA detection. Cell permeabilization and protease digestion were performed by incubating slides with 1 µg/mL Proteinase K in 0.5% PBST (Tween-20) at 40°C for 15 minutes. RNA profiling then proceeded according to the standard CosMx WTx 18,935-gene whole-transcriptome panel protocol, enabling subcellular-resolution spatial transcriptomics from the same cells previously imaged for protein expression.

### Protein-RNA multimodal spatial alignment

Because protein and RNA data in the SPACE-immune workflow are acquired as independent sequential imaging runs on the same slide, post-acquisition computational alignment is required to register RNA transcripts into the coordinate space of the protein imaging run and reassign them to the protein-derived cell segmentation boundaries. To accomplish this, we applied the Bruker Spatial Biology CosMx Multiomics Alignment Tool (available at https://github.com/Nanostring-Biostats/CosMx-Analysis-Scratch-Space), which implements a two-step registration workflow.

In the first step, per-field-of-view (FOV) two-dimensional translational shift vectors are computed by cross-correlating morphology images acquired during the protein and RNA runs, correcting for lateral displacement introduced by slide removal and reinsertion between the two imaging sessions. FOVs exhibiting anomalous displacement patterns – indicative of local cell loss or tissue disturbance between runs – are automatically flagged and excluded from downstream processing, with exclusion events logged for quality review. In the second step, the computed shift vectors are applied to RNA transcript coordinates, and transcripts are reassigned to the cell segmentation labels established during the protein imaging run. This produces a spatially unified multimodal data object in which each cell carries both its protein expression profile and its whole-transcriptome RNA measurements, anchored to a common cellular coordinate system derived from the protein run’s segmentation.

### Naïve CD4+ T cell subclustering and trajectory analysis

To investigate transcriptional heterogeneity within the naïve CD4 T-cell compartment, cells were first isolated in silico by protein-based gating, selecting cells with high CCR7 and CD45RA protein expression co-occurring with CD3 and CD4 positivity – a surface phenotype canonical for circulating naïve CD4 T-cell. This protein-anchored gating strategy was deliberately chosen over RNA-based filtering to avoid excluding cells that had undergone transcriptional reprogramming while retaining their naïve surface protein phenotype, a dissociation that itself represents a central biological finding of this study.

Transcriptomic subclustering of protein-gated naïve CD4 T cells was performed using Leiden clustering applied to a neighborhood graph derived from the whole-transcriptome RNA data. Four primary subclusters were annotated based on their marker gene profiles and named: Naïve-resting, Effector-primed, Expansion-ready, and IFN-stimulated.

Pseudotime trajectory analysis was performed using Monocle3 to order cells along a continuous axis of transcriptional progression [19, 20], with the Naïve-resting cluster designated as the trajectory root based on its marker gene profile and lowest transcriptional divergence from the canonical naïve state. Trajectory learning was performed on UMAP coordinates. Pseudotime values revealed that the Expansion-ready cluster occupied the greatest transcriptional distance from the root (mean pseudotime 7.26), indicating that it represents a state of deep metabolic and transcriptional reprogramming rather than an early transitional phenotype – a reinterpretation supported by its marker gene profile. The IFN-stimulated cluster was identified as a donor-stratified subpopulation present in approximately half of donors, consistent with bystander IFN-γ stimulation from innate cells rather than direct cytokine engagement and corroborated by elevated CXCL9 and CXCL11 in the supernatant proteomics data from the same donors.

### Supernatant cytokine profiling

Cell culture supernatants from 12 paired donors were analyzed by a colocalization-by-linkage-based multiplex proteomics platform (nELISA, Nomic Bio). Donors were stratified into IFN-stimulated cell-enriched (n = 6) and non-enriched (n = 6) groups based on IFN-stimulated cluster proportion. Between-group comparisons were performed using Wilcoxon rank-sum tests.

### HieraType multimodal cell typing

Hierarchical cell type annotation was performed using HieraType (available at https://github.com/Nanostring-Biostats/CosMx-Analysis-Scratch-Space), a fully supervised, marker-guided classification framework that jointly leverages RNA and protein measurements to assign probabilistic cell type labels. For each cell type, classification is driven by curated marker lists specifying both RNA and protein predictors, enabling HieraType to exploit the complementary information content of both modalities simultaneously rather than performing sequential or modality-specific classification. Internally, HieraType constructs continuous per-cell marker metagene scores and probabilistically assigns cell type identity via Gaussian mixture models, producing a posterior probability distribution over all candidate cell types for each cell.

Prior to classification, protein measurements were normalized to account for background signal and count distribution structure, and a joint low-dimensional embedding was derived from the combined RNA–protein covariance structure. A cell– cell similarity graph constructed from this joint embedding was passed to HieraType as a neighborhood adjacency matrix, enabling spatially coherent label propagation across transcriptomically and proteomically similar cells.

Classification followed a multi-level hierarchical pipeline mirroring the biological organization of PBMC lineages. At the broadest level, major immune lineages were identified jointly from RNA and protein markers. T cell identity was then resolved at an intermediate level distinguishing CD4 and CD8 lineages, followed by fine-grained subtype classification of CD4 T cell subtypes, CD8 T cell subtypes, and B cell subtypes. Each classification level was conditioned on posterior probabilities from the parent level, ensuring that granular subtype calls were restricted to cells for which broad lineage identity had already been established with sufficient confidence. Final cell type labels were assigned as the highest-posterior-probability call at the most granular resolved level, yielding a 17-subtype taxonomy across 4 major PBMC lineages.

### PBMC compositional analysis and Treg:Th17 ratio evaluation

Following HieraType cell type assignment, PBMC composition was quantified as the percentage of each annotated cell type out of total profiled cells per sample, rather than absolute cell counts, as total cell numbers across CosMx SMI samples reflect imaging depth and field-of-view selection rather than biological expansion. Compositional changes between treated and control conditions were assessed across all 17 annotated subtypes using paired Wilcoxon signed-rank tests across the 12 donor pairs, with exact p-values suppressed due to ties in count data.

The Treg:Th17 ratio was computed per donor as the percentage of CD4 Treg cells divided by the percentage of CD4 Th17 cells in each condition, and the paired shift in this ratio between control and treated was tested using a paired Wilcoxon signed-rank test. This ratio was chosen as the primary compositional summary statistic for the CD4 effector balance because it captures the directional relationship between two functionally antagonistic lineages in a single metric, amplifying the signal of coordinated compositional remodeling that might be missed by analyzing each subtype independently.

Differential expression between treated and control conditions was performed separately for CD4 Treg and CD4 Th17 cells using a Wilcoxon rank-sum test at the single-cell level, with genes pre-filtered by minimum detection frequency and fold-change prior to testing. Multiple comparisons were corrected using the Benjamini– Hochberg procedure. Volcano plots were generated using the EnhancedVolcano R package with pre-specified significance thresholds for adjusted p-value and log2 fold-change. Results were interpreted in conjunction with the paired compositional analysis, with particular attention to dissociations between compositional and transcriptional responses – most notably the paradoxical downregulation of IL17A in Th17 cells despite their compositional expansion, interpreted as post-activation transcriptional quiescence at the 48-hour timepoint.

### NF-κB p65 nuclear-to-cytoplasmic ratio analysis

A defining capability of imaging-based suspension cell profiling is the ability to resolve the subcellular distribution of molecules within individual cells – a readout inaccessible to conventional sequencing-based PBMC technologies. We exploited this capability to develop a nuclear-to-cytoplasmic ratio (NCR) framework for quantifying NF-κB pathway activation state at single-cell resolution, integrating both protein localization and RNA compartmentalization as orthogonal readouts of the same signaling event.

NF-κB p65 protein activation was quantified from per-cell nuclear and cytoplasmic average fluorescence intensities measured during the protein imaging run, from which the nuclear-to-cytoplasmic intensity ratio was computed for each cell. Activated cells were identified using a two-gate hierarchical strategy: a minimum mean p65 fluorescence intensity threshold to exclude p65-negative or p65-low cells, followed by an NCR threshold to confirm nuclear enrichment above resting-state levels. This biaxial gating approach was preferred over single-metric thresholding because it separates cells with genuinely elevated and nuclear-enriched p65 from those with non-specific or uniformly low signal, reducing false-positive activation calls.

To extend the NCR concept to the transcriptomic modality, we leveraged the CosMx SMI platform’s subcellular transcript assignment – which localizes each detected RNA molecule to either the nuclear or cytoplasmic compartment of its parent cell via DAPI-defined nuclear segmentation – to compute a per-gene NCR fold-change comparing NF-κB–activated versus baseline cells within each cell type. Nuclear shift was defined as NCR FC > 1.5, cytoplasmic shift as NCR FC < 0.67, and unchanged as the interval 0.67–1.5. The proportion of genes falling into each category was computed per cell type, and cell types were grouped into three relocalization profiles: minimal relocalization, in which p65 nuclear translocation is accompanied by few nuclear-or cytoplasmic-shifted genes; unidirectional relocalization, in which shifted genes are predominantly cytoplasmic; and bidirectional relocalization, in which substantial fractions of both nuclear- and cytoplasmic-shifted genes are present. This framework was applied across all 17 annotated PBMC subtypes, enabling systematic comparison of NF-κB signal propagation fidelity across immune lineages.

## Supporting Information

**Figure S1. High-quality PBMC array made on CosMx compatible slide surface with optimized procedure**

**Figure S2. Primary immune lineage cells detected and per-donor cell distribution on UMAP**

**Figure S3. Single-cell immuno-gating enabled by SPACE-immune protein data**

**Figure S4. SPACE-immune revealed two distinct activation pathways for naïve CD4 T cells upon cytokine treatment**

**Figure S5. Age-dependent immunosenescence noted as the composition of adaptive immune and innate immune cells shifted**

**Figure S6. Differential expression analysis for CD4_Th1 cells upon cytokine treatment**

**Table S1. Core immune subtyping enabled by the CosMx 64-plex IO protein panel**

**Table S2. Summary of immune composition (absolute cell number) per donor**

**Table S3. Summary of immune composition (percentage of cells) per donor**

## References

1. Nakaya, H.I., et al., Systems biology of vaccination for seasonal influenza in humans. Nat Immunol, 2011. 12(8): p. 786–95.

2. Perez, R.K., et al., Single-cell RNA-seq reveals cell type-specific molecular and genetic associations to lupus. Science, 2022. 376(6589): p. eabf1970.

3. Zhang, B., et al., Multimodal single-cell datasets characterize antigen-specific CD8. Nat Immunol, 2023. 24(10): p. 1725–1734.

4. Shimagami, H., et al., Single-cell analysis reveals immune cell abnormalities underlying the clinical heterogeneity of patients with systemic sclerosis. Nat Commun, 2025. 16(1): p. 4949.

5. Dyikanov, D., et al., Comprehensive peripheral blood immunoprofiling reveals five immunotypes with immunotherapy response characteristics in patients with cancer. Cancer Cell, 2024. 42(5): p. 759–779.e12.

6. Macosko, E.Z., et al., Highly Parallel Genome-wide Expression Profiling of Individual Cells Using Nanoliter Droplets. Cell, 2015. 161(5): p. 1202–1214.

7. Stoeckius, M., et al., Simultaneous epitope and transcriptome measurement in single cells. Nat Methods, 2017. 14(9): p. 865–868.

8. Spitzer, M.H. and G.P. Nolan, Mass Cytometry: Single Cells, Many Features. Cell, 2016. 165(4): p. 780–91.

9. Basiji, D.A., et al., Cellular image analysis and imaging by flow cytometry. Clin Lab Med, 2007. 27(3): p. 653–70, viii.

10. McGuire, D., Integrated RNA + protein multiomic cell typing using HieraType, in CosMx Analysis Scratch Space. 2025.

11. He, S., et al., High-plex imaging of RNA and proteins at subcellular resolution in fixed tissue by spatial molecular imaging. Nat Biotechnol, 2022. 40(12): p. 1794–1806.

12. Khafizov, R., et al., Sub-cellular imaging of the entire protein-coding human transcriptome (18933-plex) on FFPE tissue using spatial molecular imaging. bioRxiv, 2024.

13. Högel-Starck, C., et al., Image-based profiling of healthy donor immune cells with Blood Cell Painting reveals novel genotype-phenotype associations. bioRxiv, 2025: p. 2024.05.17.594648.

14. Pitino, E., et al., STAMP: Single-cell transcriptomics analysis and multimodal profiling through imaging. Cell, 2025. 188(18): p. 5100–5117.e26.

15. Sallusto, F., J. Geginat, and A. Lanzavecchia, Central memory and effector memory T cell subsets: function, generation, and maintenance. Annu Rev Immunol, 2004. 22: p. 745–63.

16. Filippov, I., L. Schauser, and P. Peterson, An integrated single-cell atlas of blood immune cells in aging. NPJ Aging, 2024. 10(1): p. 59.

17. Terekhova, M., P. Bohacova, and M.N. Artyomov, Human immune aging. Immunity, 2025. 58(11): p. 2646–2669.

18. Wang, Y., et al., Integrating single-cell RNA and T cell/B cell receptor sequencing with mass cytometry reveals dynamic trajectories of human peripheral immune cells from birth to old age. Nat Immunol, 2025. 26(2): p. 308–322.

19. Trapnell, C., et al., The dynamics and regulators of cell fate decisions are revealed by pseudotemporal ordering of single cells. Nat Biotechnol, 2014. 32(4): p. 381–386.

20. Cao, J., et al., The single-cell transcriptional landscape of mammalian organogenesis. Nature, 2019. 566(7745): p. 496–502.

