## Supporting Information for "Subcellular multiomic profiling with SPACE-immune reveals divergent immune activation states and intracellular signaling of PBMCs"

### Slide 1
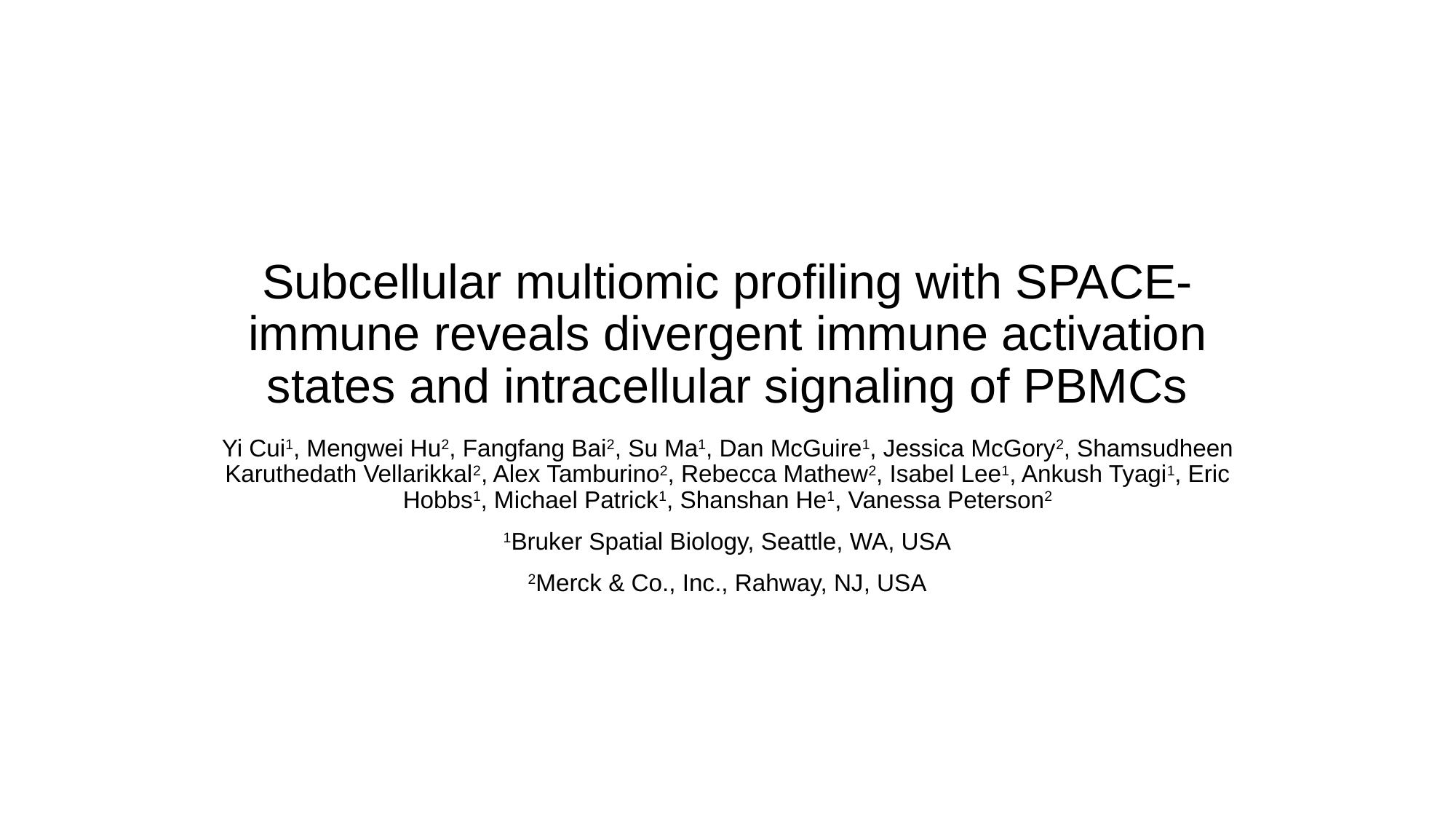

#### Subcellular multiomic profiling with SPACE-immune reveals divergent immune activation states and intracellular signaling of PBMCs
Yi Cui^1^, Mengwei Hu^2^, Fangfang Bai^2^, Su Ma^1^, Dan McGuire^1^, Jessica McGory^2^, Shamsudheen Karuthedath Vellarikkal^2^, Alex Tamburino^2^, Rebecca Mathew^2^, Isabel Lee^1^, Ankush Tyagi^1^, Eric Hobbs^1^, Michael Patrick^1^, Shanshan He^1^, Vanessa Peterson^2^
^1^Bruker Spatial Biology, Seattle, WA, USA
^2^Merck & Co., Inc., Rahway, NJ, USA

### Slide 2
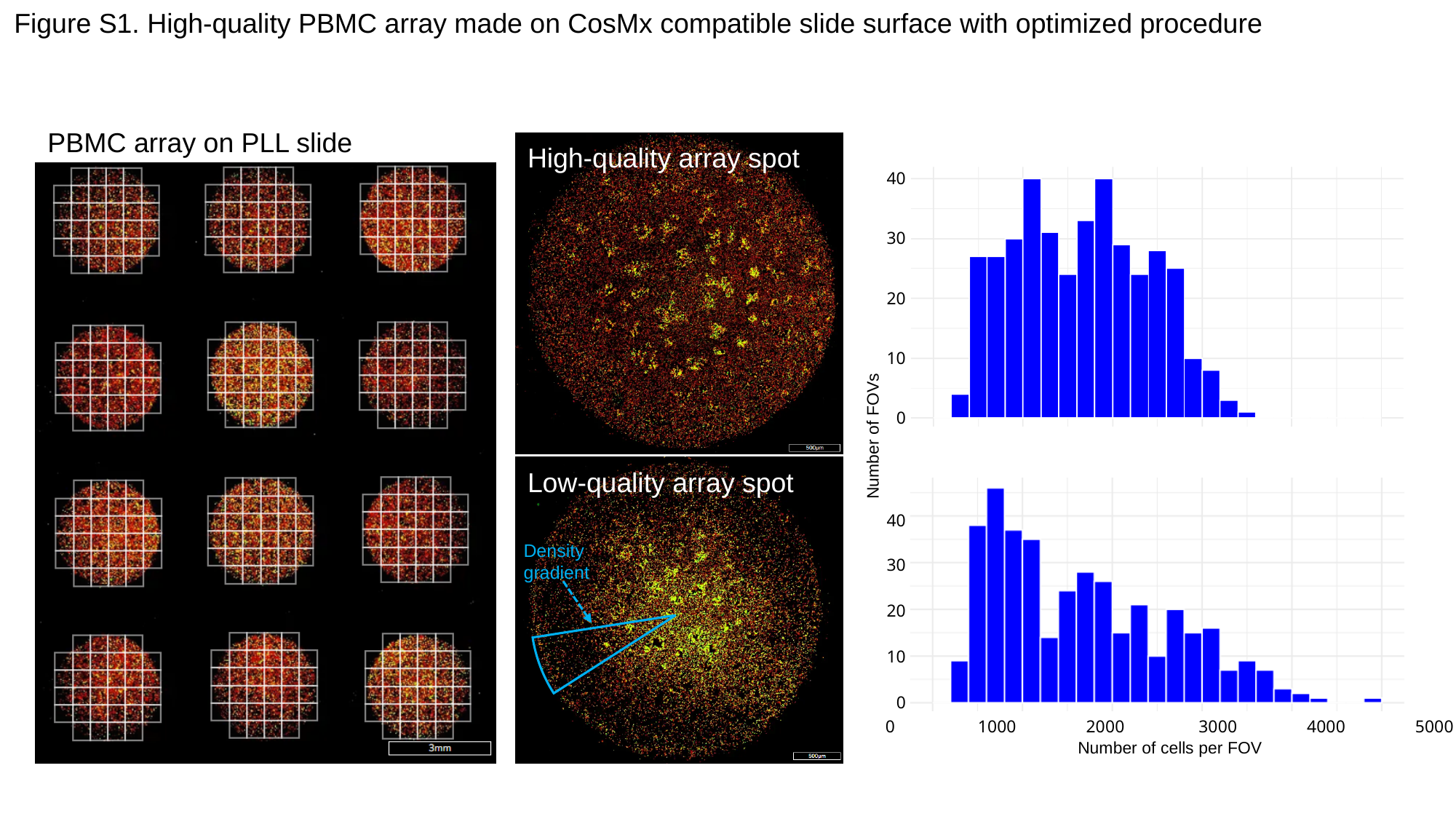

Figure S1. High-quality PBMC array made on CosMx compatible slide surface with optimized procedure
PBMC array on PLL slide
High-quality array spot
40
30
20
10
0
Number of FOVs
Density gradient
Low-quality array spot
40
30
20
10
0
0 1000 2000 3000 4000 5000
Number of cells per FOV

### Slide 3
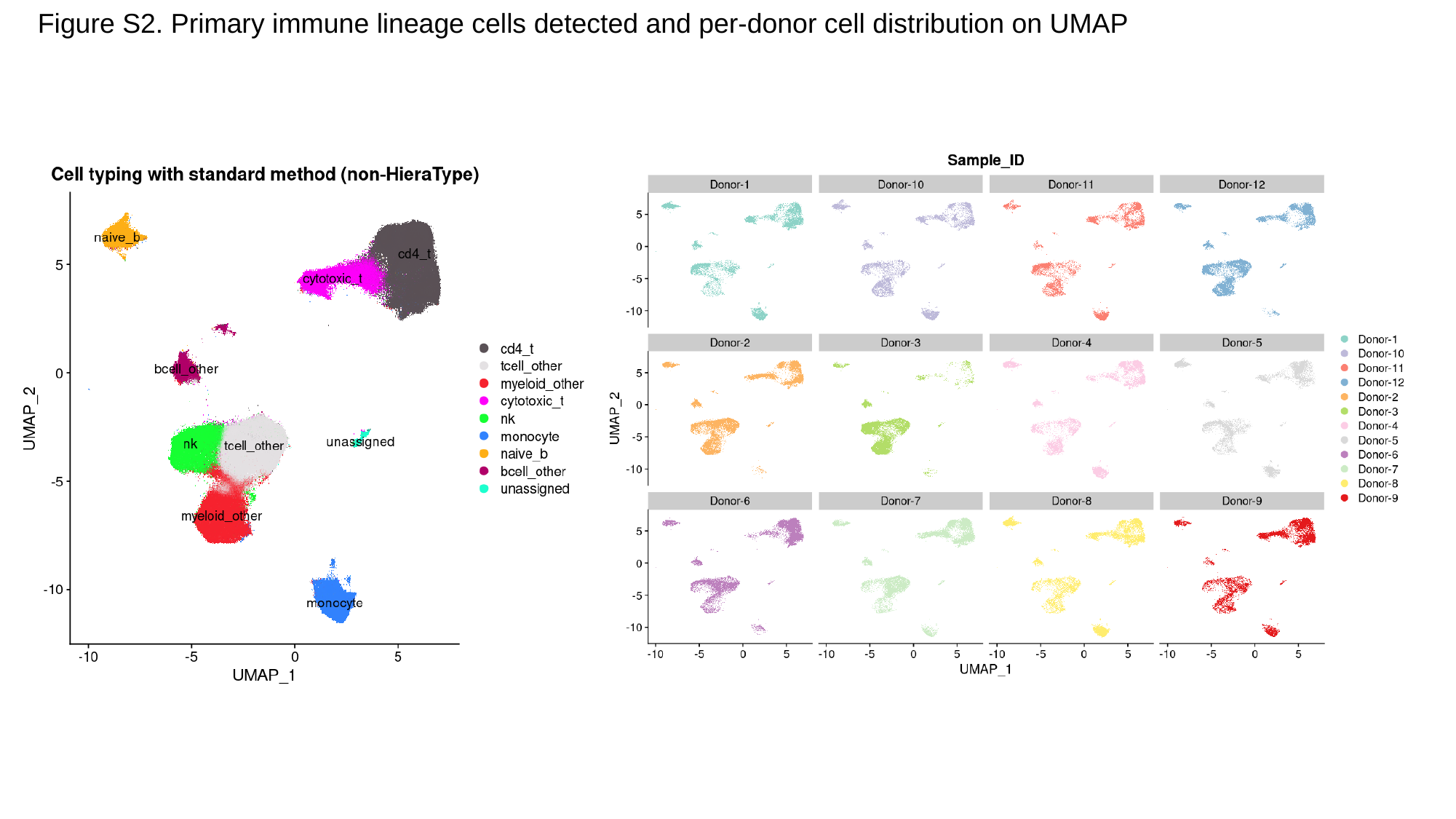

Figure S2. Primary immune lineage cells detected and per-donor cell distribution on UMAP

### Slide 4
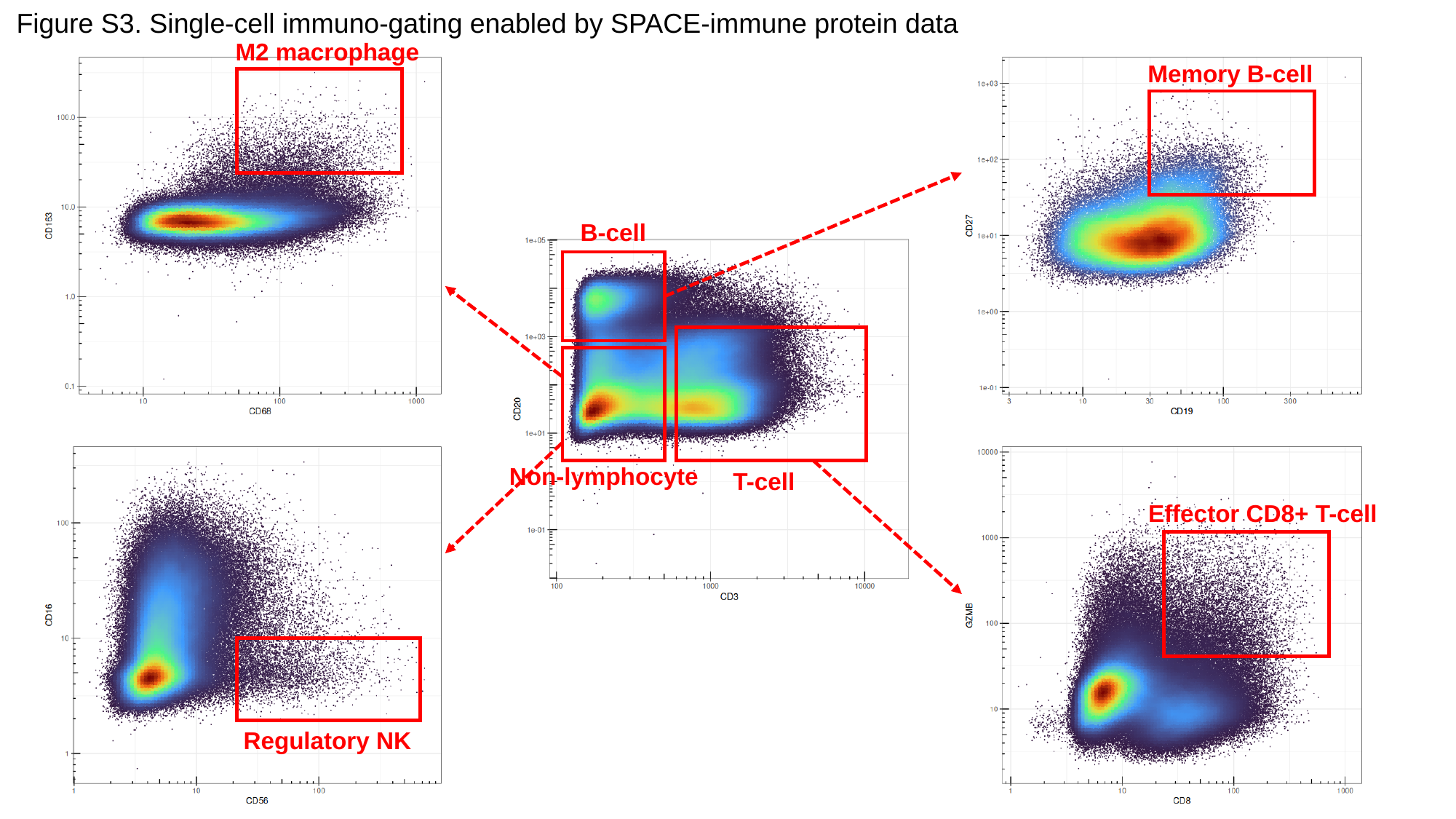

Figure S3. Single-cell immuno-gating enabled by SPACE-immune protein data
M2 macrophage
Memory B-cell
B-cell
Non-lymphocyte
T-cell
Effector CD8+ T-cell
Regulatory NK

### Slide 5
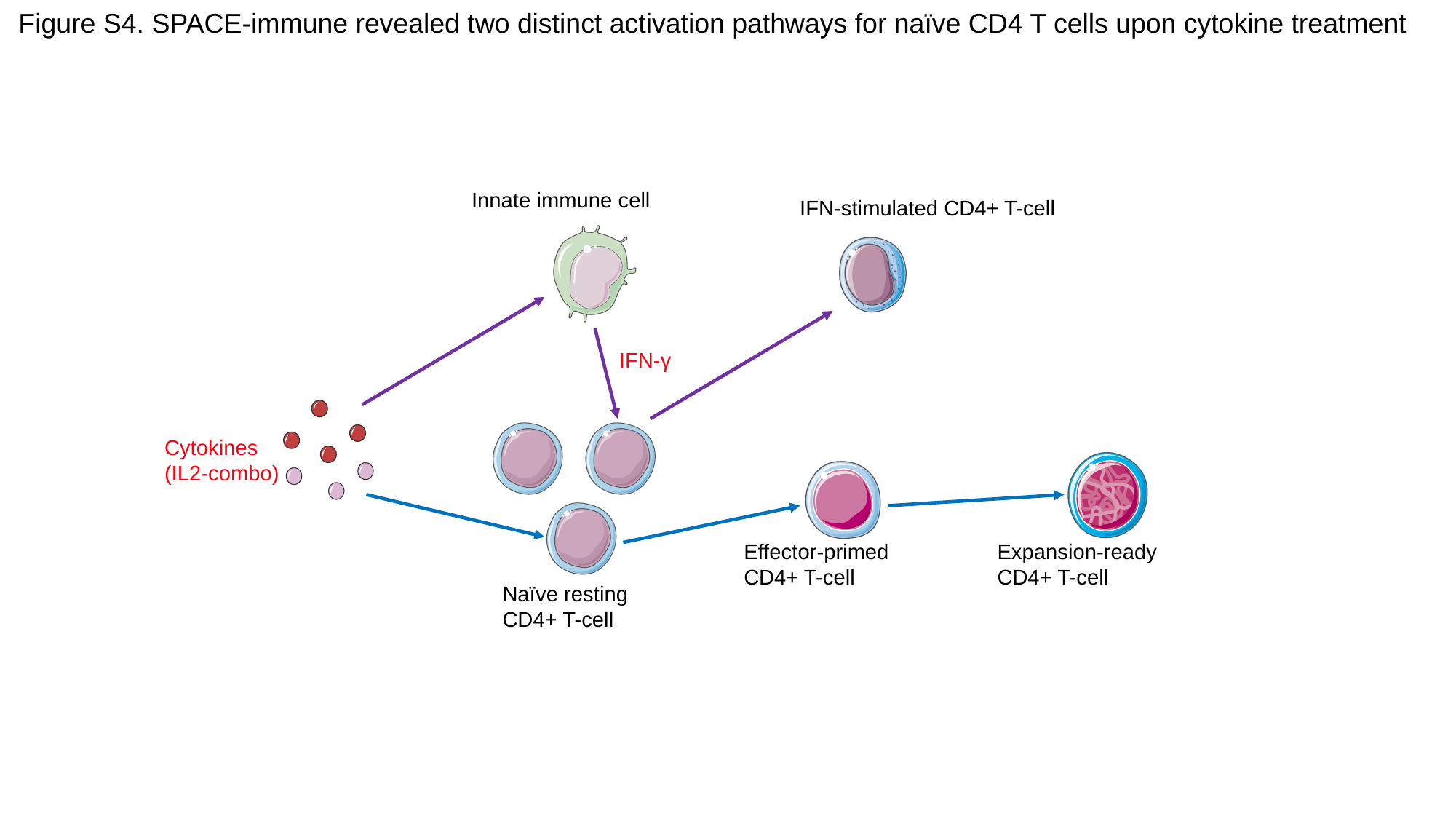

Figure S4. SPACE-immune revealed two distinct activation pathways for naïve CD4 T cells upon cytokine treatment
Innate immune cell
IFN-stimulated CD4+ T-cell
IFN-γ
Cytokines
(IL2-combo)
Effector-primed
CD4+ T-cell
Expansion-ready
CD4+ T-cell
Naïve resting
CD4+ T-cell

### Slide 6
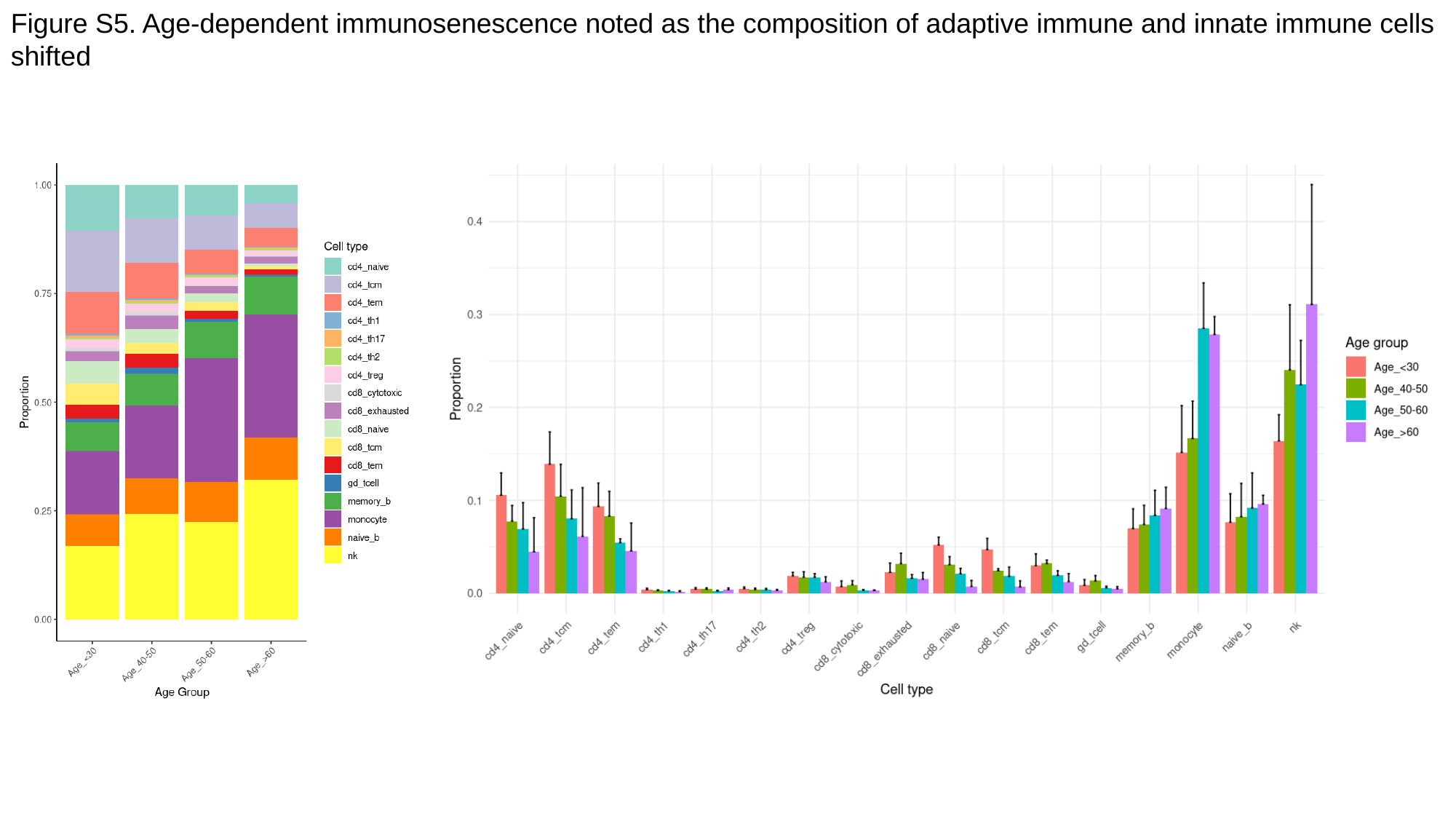

Figure S5. Age-dependent immunosenescence noted as the composition of adaptive immune and innate immune cells shifted

### Slide 7
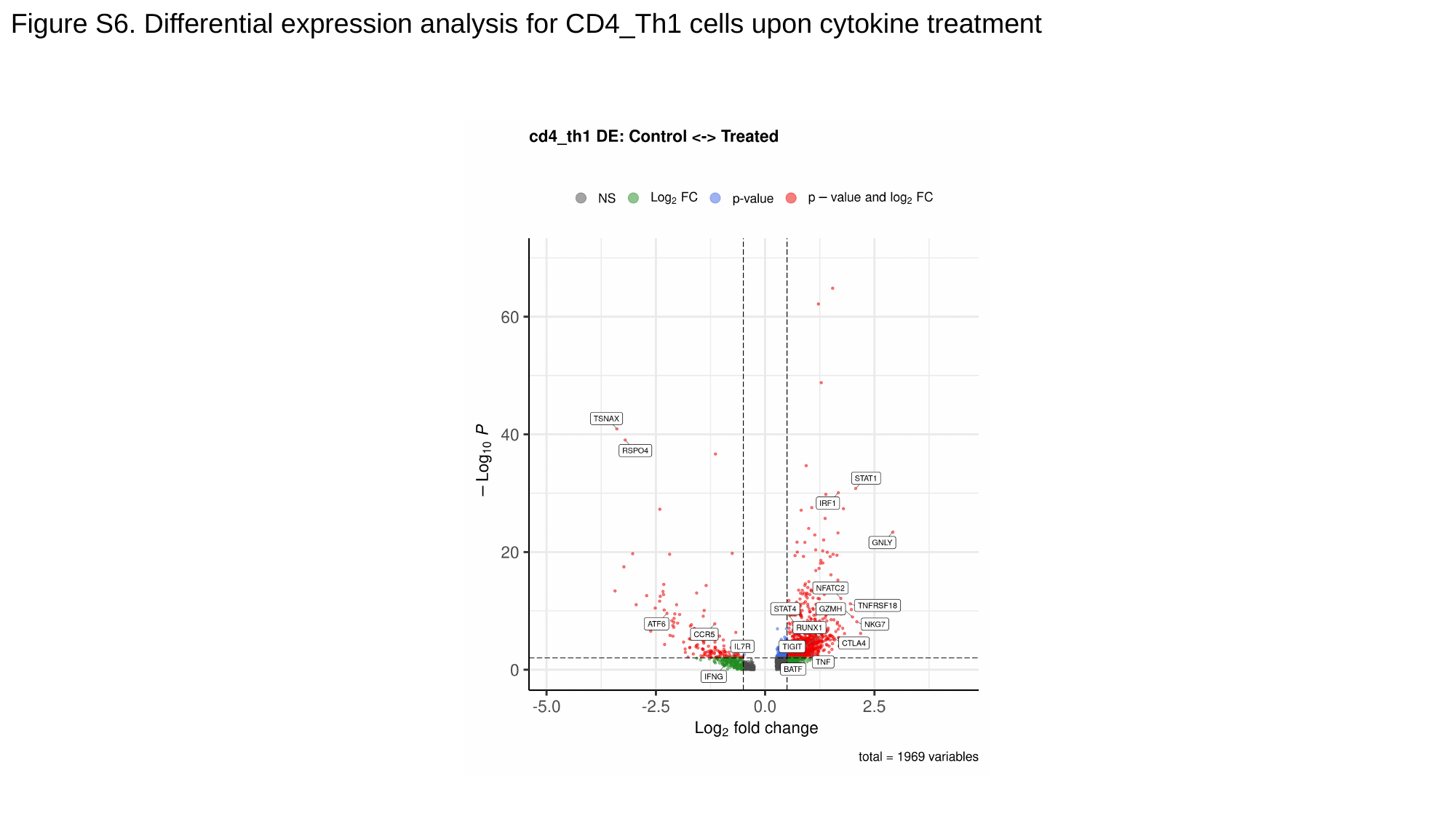

Figure S6. Differential expression analysis for CD4_Th1 cells upon cytokine treatment

### Slide 8
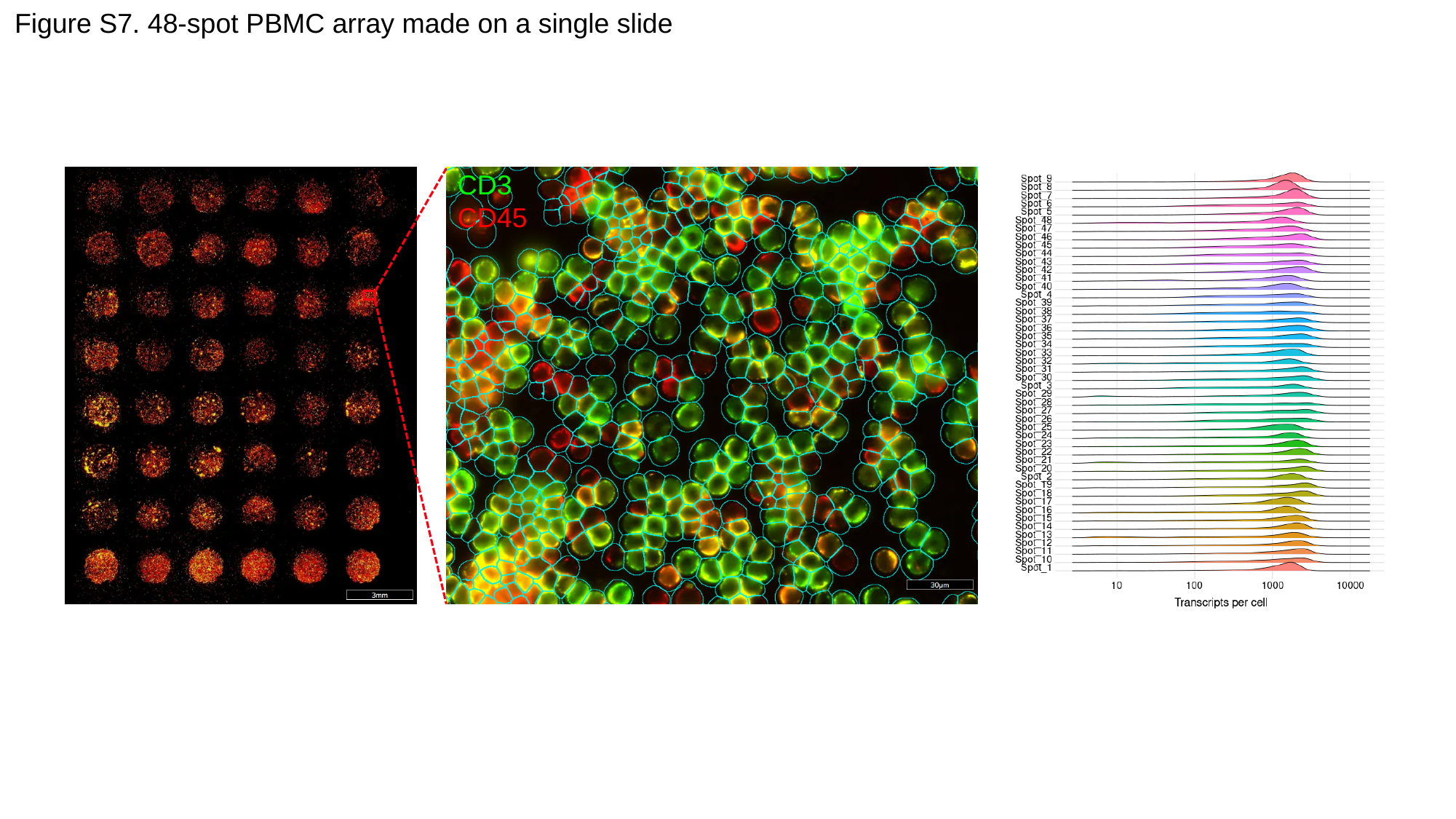

Figure S7. 48-spot PBMC array made on a single slide
CD3
CD45

### Slide 9
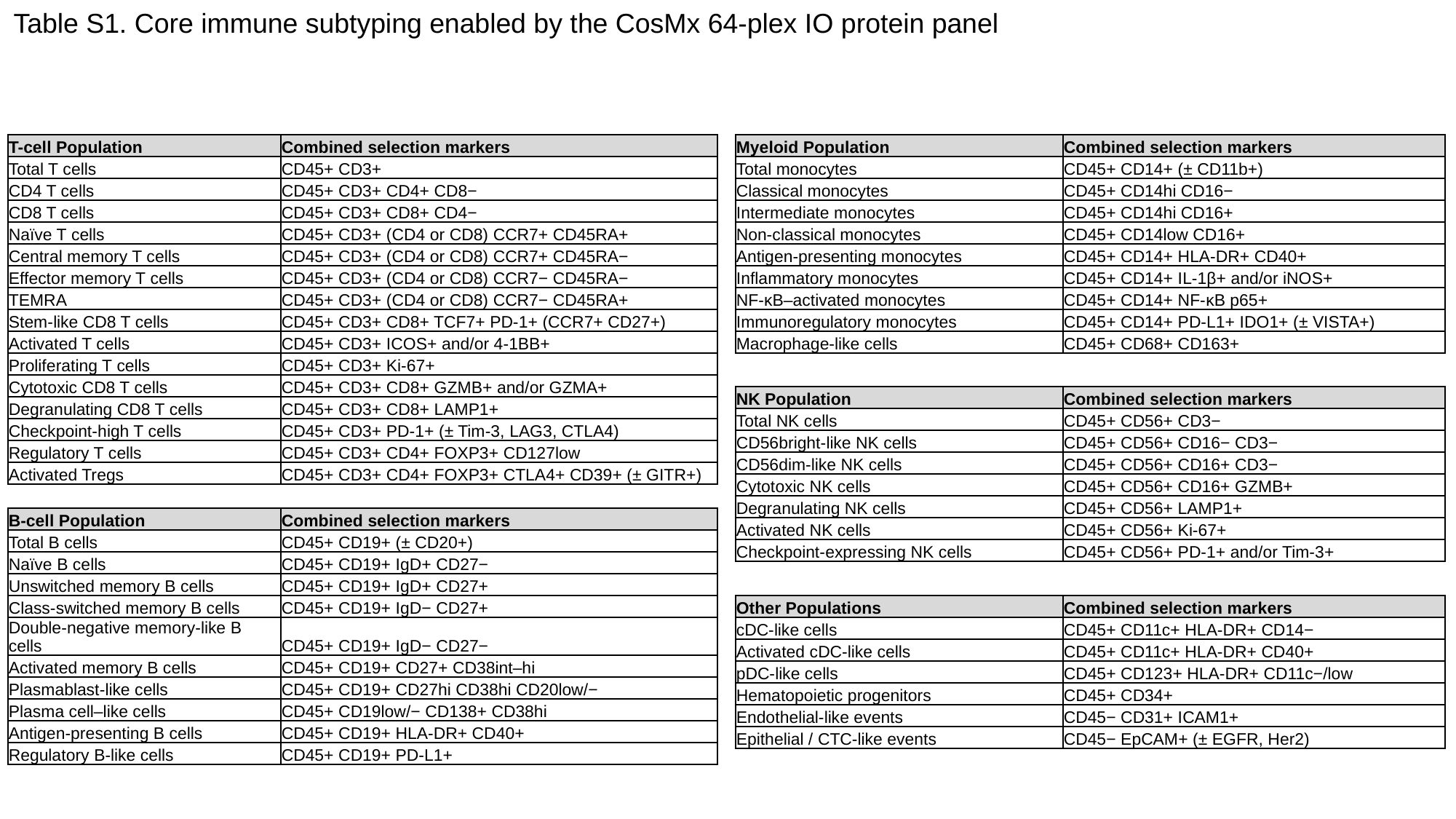

Table S1. Core immune subtyping enabled by the CosMx 64-plex IO protein panel
| T-cell Population | Combined selection markers |
| --- | --- |
| Total T cells | CD45+ CD3+ |
| CD4 T cells | CD45+ CD3+ CD4+ CD8− |
| CD8 T cells | CD45+ CD3+ CD8+ CD4− |
| Naïve T cells | CD45+ CD3+ (CD4 or CD8) CCR7+ CD45RA+ |
| Central memory T cells | CD45+ CD3+ (CD4 or CD8) CCR7+ CD45RA− |
| Effector memory T cells | CD45+ CD3+ (CD4 or CD8) CCR7− CD45RA− |
| TEMRA | CD45+ CD3+ (CD4 or CD8) CCR7− CD45RA+ |
| Stem-like CD8 T cells | CD45+ CD3+ CD8+ TCF7+ PD-1+ (CCR7+ CD27+) |
| Activated T cells | CD45+ CD3+ ICOS+ and/or 4-1BB+ |
| Proliferating T cells | CD45+ CD3+ Ki-67+ |
| Cytotoxic CD8 T cells | CD45+ CD3+ CD8+ GZMB+ and/or GZMA+ |
| Degranulating CD8 T cells | CD45+ CD3+ CD8+ LAMP1+ |
| Checkpoint-high T cells | CD45+ CD3+ PD-1+ (± Tim-3, LAG3, CTLA4) |
| Regulatory T cells | CD45+ CD3+ CD4+ FOXP3+ CD127low |
| Activated Tregs | CD45+ CD3+ CD4+ FOXP3+ CTLA4+ CD39+ (± GITR+) |
| Myeloid Population | Combined selection markers |
| --- | --- |
| Total monocytes | CD45+ CD14+ (± CD11b+) |
| Classical monocytes | CD45+ CD14hi CD16− |
| Intermediate monocytes | CD45+ CD14hi CD16+ |
| Non-classical monocytes | CD45+ CD14low CD16+ |
| Antigen-presenting monocytes | CD45+ CD14+ HLA-DR+ CD40+ |
| Inflammatory monocytes | CD45+ CD14+ IL-1β+ and/or iNOS+ |
| NF-κB–activated monocytes | CD45+ CD14+ NF-κB p65+ |
| Immunoregulatory monocytes | CD45+ CD14+ PD-L1+ IDO1+ (± VISTA+) |
| Macrophage-like cells | CD45+ CD68+ CD163+ |
| NK Population | Combined selection markers |
| --- | --- |
| Total NK cells | CD45+ CD56+ CD3− |
| CD56bright-like NK cells | CD45+ CD56+ CD16− CD3− |
| CD56dim-like NK cells | CD45+ CD56+ CD16+ CD3− |
| Cytotoxic NK cells | CD45+ CD56+ CD16+ GZMB+ |
| Degranulating NK cells | CD45+ CD56+ LAMP1+ |
| Activated NK cells | CD45+ CD56+ Ki-67+ |
| Checkpoint-expressing NK cells | CD45+ CD56+ PD-1+ and/or Tim-3+ |
| B-cell Population | Combined selection markers |
| --- | --- |
| Total B cells | CD45+ CD19+ (± CD20+) |
| Naïve B cells | CD45+ CD19+ IgD+ CD27− |
| Unswitched memory B cells | CD45+ CD19+ IgD+ CD27+ |
| Class-switched memory B cells | CD45+ CD19+ IgD− CD27+ |
| Double-negative memory-like B cells | CD45+ CD19+ IgD− CD27− |
| Activated memory B cells | CD45+ CD19+ CD27+ CD38int–hi |
| Plasmablast-like cells | CD45+ CD19+ CD27hi CD38hi CD20low/− |
| Plasma cell–like cells | CD45+ CD19low/− CD138+ CD38hi |
| Antigen-presenting B cells | CD45+ CD19+ HLA-DR+ CD40+ |
| Regulatory B-like cells | CD45+ CD19+ PD-L1+ |
| Other Populations | Combined selection markers |
| --- | --- |
| cDC-like cells | CD45+ CD11c+ HLA-DR+ CD14− |
| Activated cDC-like cells | CD45+ CD11c+ HLA-DR+ CD40+ |
| pDC-like cells | CD45+ CD123+ HLA-DR+ CD11c−/low |
| Hematopoietic progenitors | CD45+ CD34+ |
| Endothelial-like events | CD45− CD31+ ICAM1+ |
| Epithelial / CTC-like events | CD45− EpCAM+ (± EGFR, Her2) |

### Slide 10
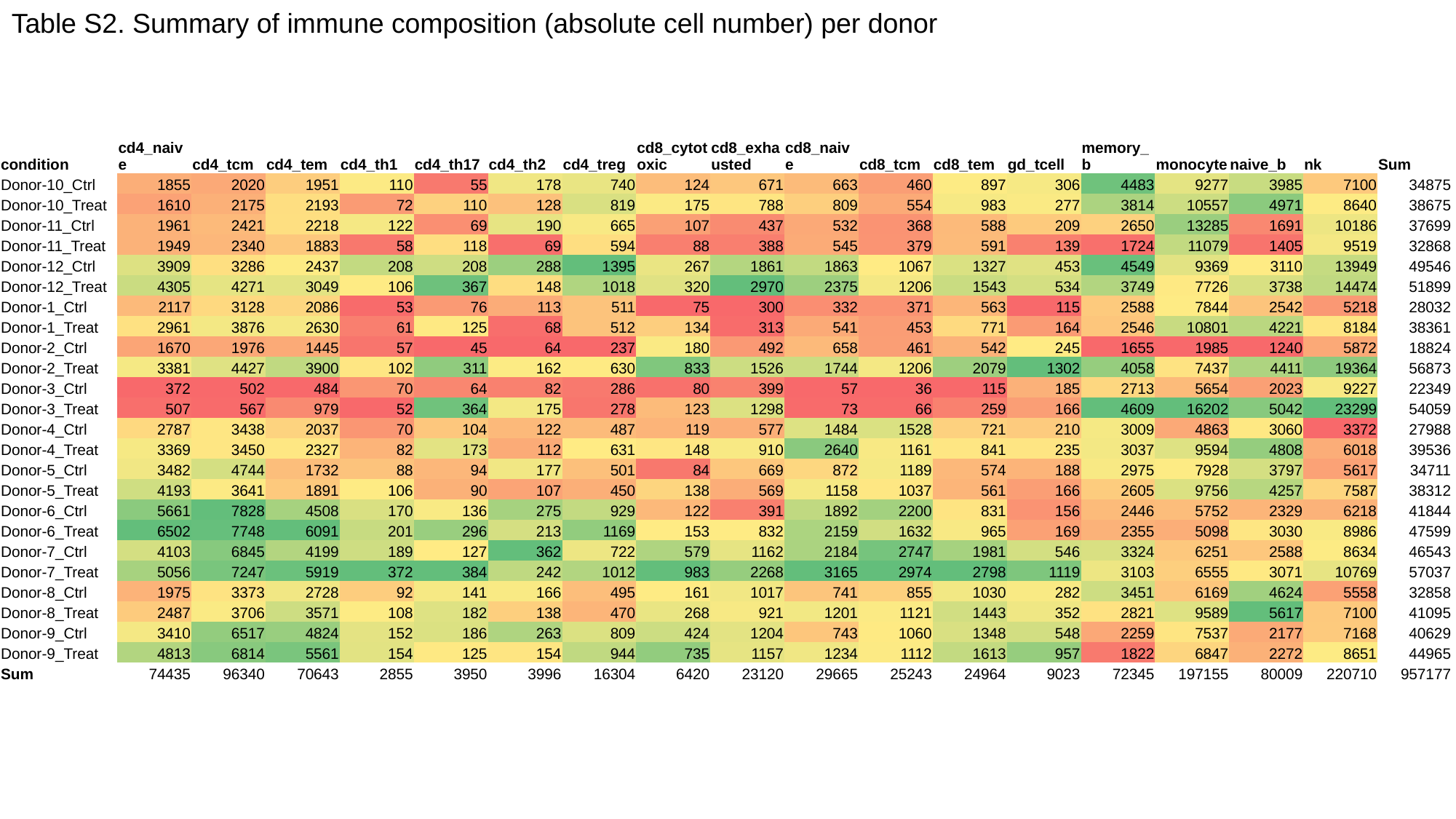

Table S2. Summary of immune composition (absolute cell number) per donor
| condition | cd4\_naive | cd4\_tcm | cd4\_tem | cd4\_th1 | cd4\_th17 | cd4\_th2 | cd4\_treg | cd8\_cytotoxic | cd8\_exhausted | cd8\_naive | cd8\_tcm | cd8\_tem | gd\_tcell | memory\_b | monocyte | naive\_b | nk | Sum |
| --- | --- | --- | --- | --- | --- | --- | --- | --- | --- | --- | --- | --- | --- | --- | --- | --- | --- | --- |
| Donor-10\_Ctrl | 1855 | 2020 | 1951 | 110 | 55 | 178 | 740 | 124 | 671 | 663 | 460 | 897 | 306 | 4483 | 9277 | 3985 | 7100 | 34875 |
| Donor-10\_Treat | 1610 | 2175 | 2193 | 72 | 110 | 128 | 819 | 175 | 788 | 809 | 554 | 983 | 277 | 3814 | 10557 | 4971 | 8640 | 38675 |
| Donor-11\_Ctrl | 1961 | 2421 | 2218 | 122 | 69 | 190 | 665 | 107 | 437 | 532 | 368 | 588 | 209 | 2650 | 13285 | 1691 | 10186 | 37699 |
| Donor-11\_Treat | 1949 | 2340 | 1883 | 58 | 118 | 69 | 594 | 88 | 388 | 545 | 379 | 591 | 139 | 1724 | 11079 | 1405 | 9519 | 32868 |
| Donor-12\_Ctrl | 3909 | 3286 | 2437 | 208 | 208 | 288 | 1395 | 267 | 1861 | 1863 | 1067 | 1327 | 453 | 4549 | 9369 | 3110 | 13949 | 49546 |
| Donor-12\_Treat | 4305 | 4271 | 3049 | 106 | 367 | 148 | 1018 | 320 | 2970 | 2375 | 1206 | 1543 | 534 | 3749 | 7726 | 3738 | 14474 | 51899 |
| Donor-1\_Ctrl | 2117 | 3128 | 2086 | 53 | 76 | 113 | 511 | 75 | 300 | 332 | 371 | 563 | 115 | 2588 | 7844 | 2542 | 5218 | 28032 |
| Donor-1\_Treat | 2961 | 3876 | 2630 | 61 | 125 | 68 | 512 | 134 | 313 | 541 | 453 | 771 | 164 | 2546 | 10801 | 4221 | 8184 | 38361 |
| Donor-2\_Ctrl | 1670 | 1976 | 1445 | 57 | 45 | 64 | 237 | 180 | 492 | 658 | 461 | 542 | 245 | 1655 | 1985 | 1240 | 5872 | 18824 |
| Donor-2\_Treat | 3381 | 4427 | 3900 | 102 | 311 | 162 | 630 | 833 | 1526 | 1744 | 1206 | 2079 | 1302 | 4058 | 7437 | 4411 | 19364 | 56873 |
| Donor-3\_Ctrl | 372 | 502 | 484 | 70 | 64 | 82 | 286 | 80 | 399 | 57 | 36 | 115 | 185 | 2713 | 5654 | 2023 | 9227 | 22349 |
| Donor-3\_Treat | 507 | 567 | 979 | 52 | 364 | 175 | 278 | 123 | 1298 | 73 | 66 | 259 | 166 | 4609 | 16202 | 5042 | 23299 | 54059 |
| Donor-4\_Ctrl | 2787 | 3438 | 2037 | 70 | 104 | 122 | 487 | 119 | 577 | 1484 | 1528 | 721 | 210 | 3009 | 4863 | 3060 | 3372 | 27988 |
| Donor-4\_Treat | 3369 | 3450 | 2327 | 82 | 173 | 112 | 631 | 148 | 910 | 2640 | 1161 | 841 | 235 | 3037 | 9594 | 4808 | 6018 | 39536 |
| Donor-5\_Ctrl | 3482 | 4744 | 1732 | 88 | 94 | 177 | 501 | 84 | 669 | 872 | 1189 | 574 | 188 | 2975 | 7928 | 3797 | 5617 | 34711 |
| Donor-5\_Treat | 4193 | 3641 | 1891 | 106 | 90 | 107 | 450 | 138 | 569 | 1158 | 1037 | 561 | 166 | 2605 | 9756 | 4257 | 7587 | 38312 |
| Donor-6\_Ctrl | 5661 | 7828 | 4508 | 170 | 136 | 275 | 929 | 122 | 391 | 1892 | 2200 | 831 | 156 | 2446 | 5752 | 2329 | 6218 | 41844 |
| Donor-6\_Treat | 6502 | 7748 | 6091 | 201 | 296 | 213 | 1169 | 153 | 832 | 2159 | 1632 | 965 | 169 | 2355 | 5098 | 3030 | 8986 | 47599 |
| Donor-7\_Ctrl | 4103 | 6845 | 4199 | 189 | 127 | 362 | 722 | 579 | 1162 | 2184 | 2747 | 1981 | 546 | 3324 | 6251 | 2588 | 8634 | 46543 |
| Donor-7\_Treat | 5056 | 7247 | 5919 | 372 | 384 | 242 | 1012 | 983 | 2268 | 3165 | 2974 | 2798 | 1119 | 3103 | 6555 | 3071 | 10769 | 57037 |
| Donor-8\_Ctrl | 1975 | 3373 | 2728 | 92 | 141 | 166 | 495 | 161 | 1017 | 741 | 855 | 1030 | 282 | 3451 | 6169 | 4624 | 5558 | 32858 |
| Donor-8\_Treat | 2487 | 3706 | 3571 | 108 | 182 | 138 | 470 | 268 | 921 | 1201 | 1121 | 1443 | 352 | 2821 | 9589 | 5617 | 7100 | 41095 |
| Donor-9\_Ctrl | 3410 | 6517 | 4824 | 152 | 186 | 263 | 809 | 424 | 1204 | 743 | 1060 | 1348 | 548 | 2259 | 7537 | 2177 | 7168 | 40629 |
| Donor-9\_Treat | 4813 | 6814 | 5561 | 154 | 125 | 154 | 944 | 735 | 1157 | 1234 | 1112 | 1613 | 957 | 1822 | 6847 | 2272 | 8651 | 44965 |
| Sum | 74435 | 96340 | 70643 | 2855 | 3950 | 3996 | 16304 | 6420 | 23120 | 29665 | 25243 | 24964 | 9023 | 72345 | 197155 | 80009 | 220710 | 957177 |

### Slide 11
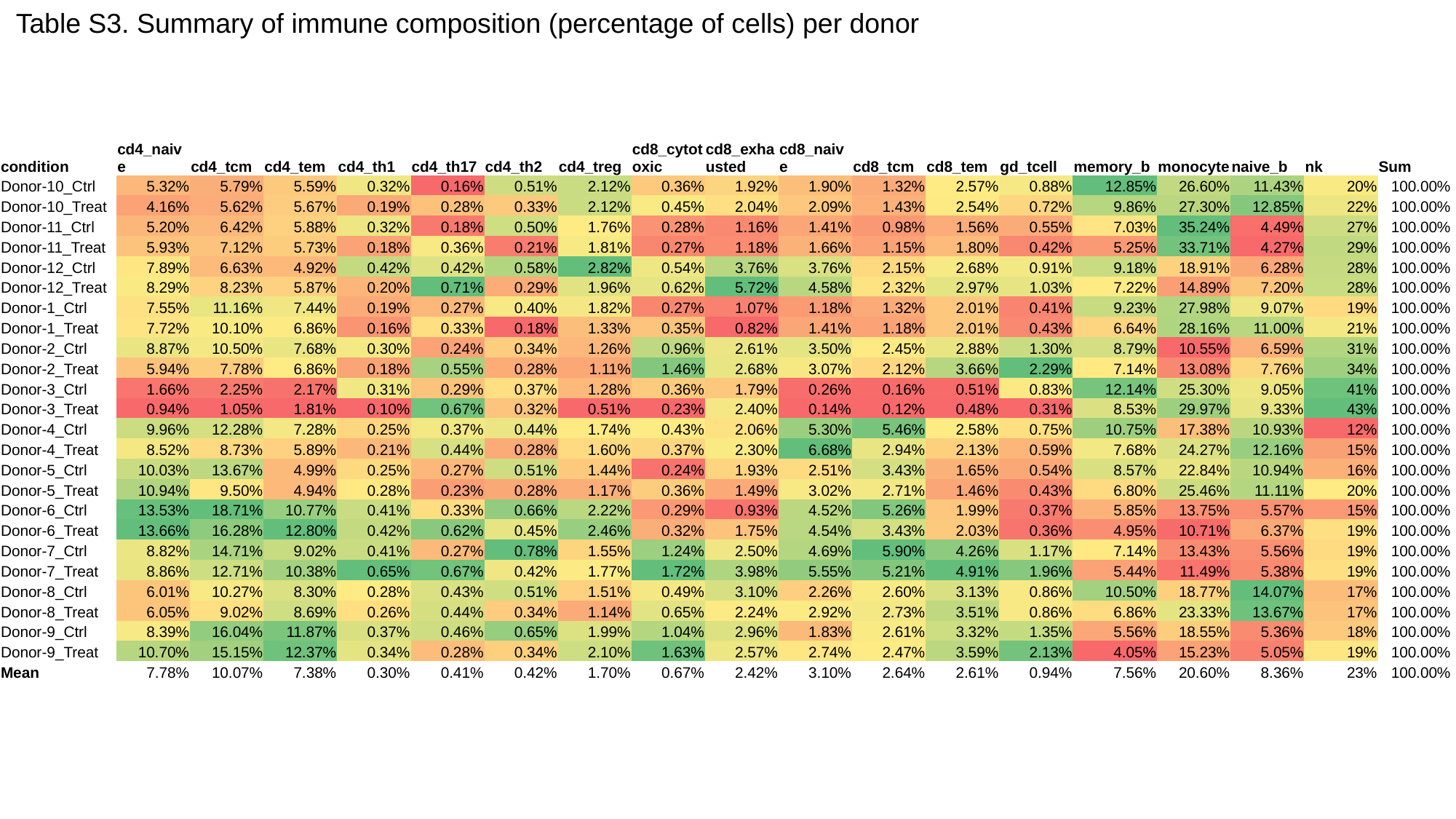

Table S3. Summary of immune composition (percentage of cells) per donor
| condition | cd4\_naive | cd4\_tcm | cd4\_tem | cd4\_th1 | cd4\_th17 | cd4\_th2 | cd4\_treg | cd8\_cytotoxic | cd8\_exhausted | cd8\_naive | cd8\_tcm | cd8\_tem | gd\_tcell | memory\_b | monocyte | naive\_b | nk | Sum |
| --- | --- | --- | --- | --- | --- | --- | --- | --- | --- | --- | --- | --- | --- | --- | --- | --- | --- | --- |
| Donor-10\_Ctrl | 5.32% | 5.79% | 5.59% | 0.32% | 0.16% | 0.51% | 2.12% | 0.36% | 1.92% | 1.90% | 1.32% | 2.57% | 0.88% | 12.85% | 26.60% | 11.43% | 20% | 100.00% |
| Donor-10\_Treat | 4.16% | 5.62% | 5.67% | 0.19% | 0.28% | 0.33% | 2.12% | 0.45% | 2.04% | 2.09% | 1.43% | 2.54% | 0.72% | 9.86% | 27.30% | 12.85% | 22% | 100.00% |
| Donor-11\_Ctrl | 5.20% | 6.42% | 5.88% | 0.32% | 0.18% | 0.50% | 1.76% | 0.28% | 1.16% | 1.41% | 0.98% | 1.56% | 0.55% | 7.03% | 35.24% | 4.49% | 27% | 100.00% |
| Donor-11\_Treat | 5.93% | 7.12% | 5.73% | 0.18% | 0.36% | 0.21% | 1.81% | 0.27% | 1.18% | 1.66% | 1.15% | 1.80% | 0.42% | 5.25% | 33.71% | 4.27% | 29% | 100.00% |
| Donor-12\_Ctrl | 7.89% | 6.63% | 4.92% | 0.42% | 0.42% | 0.58% | 2.82% | 0.54% | 3.76% | 3.76% | 2.15% | 2.68% | 0.91% | 9.18% | 18.91% | 6.28% | 28% | 100.00% |
| Donor-12\_Treat | 8.29% | 8.23% | 5.87% | 0.20% | 0.71% | 0.29% | 1.96% | 0.62% | 5.72% | 4.58% | 2.32% | 2.97% | 1.03% | 7.22% | 14.89% | 7.20% | 28% | 100.00% |
| Donor-1\_Ctrl | 7.55% | 11.16% | 7.44% | 0.19% | 0.27% | 0.40% | 1.82% | 0.27% | 1.07% | 1.18% | 1.32% | 2.01% | 0.41% | 9.23% | 27.98% | 9.07% | 19% | 100.00% |
| Donor-1\_Treat | 7.72% | 10.10% | 6.86% | 0.16% | 0.33% | 0.18% | 1.33% | 0.35% | 0.82% | 1.41% | 1.18% | 2.01% | 0.43% | 6.64% | 28.16% | 11.00% | 21% | 100.00% |
| Donor-2\_Ctrl | 8.87% | 10.50% | 7.68% | 0.30% | 0.24% | 0.34% | 1.26% | 0.96% | 2.61% | 3.50% | 2.45% | 2.88% | 1.30% | 8.79% | 10.55% | 6.59% | 31% | 100.00% |
| Donor-2\_Treat | 5.94% | 7.78% | 6.86% | 0.18% | 0.55% | 0.28% | 1.11% | 1.46% | 2.68% | 3.07% | 2.12% | 3.66% | 2.29% | 7.14% | 13.08% | 7.76% | 34% | 100.00% |
| Donor-3\_Ctrl | 1.66% | 2.25% | 2.17% | 0.31% | 0.29% | 0.37% | 1.28% | 0.36% | 1.79% | 0.26% | 0.16% | 0.51% | 0.83% | 12.14% | 25.30% | 9.05% | 41% | 100.00% |
| Donor-3\_Treat | 0.94% | 1.05% | 1.81% | 0.10% | 0.67% | 0.32% | 0.51% | 0.23% | 2.40% | 0.14% | 0.12% | 0.48% | 0.31% | 8.53% | 29.97% | 9.33% | 43% | 100.00% |
| Donor-4\_Ctrl | 9.96% | 12.28% | 7.28% | 0.25% | 0.37% | 0.44% | 1.74% | 0.43% | 2.06% | 5.30% | 5.46% | 2.58% | 0.75% | 10.75% | 17.38% | 10.93% | 12% | 100.00% |
| Donor-4\_Treat | 8.52% | 8.73% | 5.89% | 0.21% | 0.44% | 0.28% | 1.60% | 0.37% | 2.30% | 6.68% | 2.94% | 2.13% | 0.59% | 7.68% | 24.27% | 12.16% | 15% | 100.00% |
| Donor-5\_Ctrl | 10.03% | 13.67% | 4.99% | 0.25% | 0.27% | 0.51% | 1.44% | 0.24% | 1.93% | 2.51% | 3.43% | 1.65% | 0.54% | 8.57% | 22.84% | 10.94% | 16% | 100.00% |
| Donor-5\_Treat | 10.94% | 9.50% | 4.94% | 0.28% | 0.23% | 0.28% | 1.17% | 0.36% | 1.49% | 3.02% | 2.71% | 1.46% | 0.43% | 6.80% | 25.46% | 11.11% | 20% | 100.00% |
| Donor-6\_Ctrl | 13.53% | 18.71% | 10.77% | 0.41% | 0.33% | 0.66% | 2.22% | 0.29% | 0.93% | 4.52% | 5.26% | 1.99% | 0.37% | 5.85% | 13.75% | 5.57% | 15% | 100.00% |
| Donor-6\_Treat | 13.66% | 16.28% | 12.80% | 0.42% | 0.62% | 0.45% | 2.46% | 0.32% | 1.75% | 4.54% | 3.43% | 2.03% | 0.36% | 4.95% | 10.71% | 6.37% | 19% | 100.00% |
| Donor-7\_Ctrl | 8.82% | 14.71% | 9.02% | 0.41% | 0.27% | 0.78% | 1.55% | 1.24% | 2.50% | 4.69% | 5.90% | 4.26% | 1.17% | 7.14% | 13.43% | 5.56% | 19% | 100.00% |
| Donor-7\_Treat | 8.86% | 12.71% | 10.38% | 0.65% | 0.67% | 0.42% | 1.77% | 1.72% | 3.98% | 5.55% | 5.21% | 4.91% | 1.96% | 5.44% | 11.49% | 5.38% | 19% | 100.00% |
| Donor-8\_Ctrl | 6.01% | 10.27% | 8.30% | 0.28% | 0.43% | 0.51% | 1.51% | 0.49% | 3.10% | 2.26% | 2.60% | 3.13% | 0.86% | 10.50% | 18.77% | 14.07% | 17% | 100.00% |
| Donor-8\_Treat | 6.05% | 9.02% | 8.69% | 0.26% | 0.44% | 0.34% | 1.14% | 0.65% | 2.24% | 2.92% | 2.73% | 3.51% | 0.86% | 6.86% | 23.33% | 13.67% | 17% | 100.00% |
| Donor-9\_Ctrl | 8.39% | 16.04% | 11.87% | 0.37% | 0.46% | 0.65% | 1.99% | 1.04% | 2.96% | 1.83% | 2.61% | 3.32% | 1.35% | 5.56% | 18.55% | 5.36% | 18% | 100.00% |
| Donor-9\_Treat | 10.70% | 15.15% | 12.37% | 0.34% | 0.28% | 0.34% | 2.10% | 1.63% | 2.57% | 2.74% | 2.47% | 3.59% | 2.13% | 4.05% | 15.23% | 5.05% | 19% | 100.00% |
| Mean | 7.78% | 10.07% | 7.38% | 0.30% | 0.41% | 0.42% | 1.70% | 0.67% | 2.42% | 3.10% | 2.64% | 2.61% | 0.94% | 7.56% | 20.60% | 8.36% | 23% | 100.00% |
